# Comparative Proteogenomic Analysis of Right-sided Colon Cancer, Left-sided Colon Cancer and Rectal Cancers Reveal Distinct Mutational Profiles

**DOI:** 10.1101/359679

**Authors:** Robin Imperial, Zaheer Ahmed, Omer M Toor, Cihat Erdoğan, Ateeq Khaliq, Paul Case, James Case, Kevin Kennedy, Lee S. Cummings, Niklas Melton, Shahzad Raza, Banu Diri, Ramzi Mohammad, Bassel El-Rayes, Timothy Pluard, Arif Hussain, Janakiraman Subramanian, Ashiq Masood

**Author notes:** Corresponding Authors Ashiq Masood, MD Assistant Professor of Medicine Center for Precision Oncology Gastrointestinal Cancer Program Saint Luke’s Cancer Institute, University of Missouri School of Medicine, Kansas City 4321 Washington St. Kansas City, MO 64111.

## Abstract

To understand the molecular differences between right-sided colon cancer (RCC), left-sided colon cancer (LCC) and rectal cancer, we analyzed colorectal tumors at the DNA, RNA, miRNA and protein levels using previously sequenced data from The Cancer Genome Atlas and Memorial Sloan Kettering Cancer Center. Clonal evolution analysis identified the same tumor-initiating events involving APC, KRAS and TP53 genes in RCC, LCC and rectal cancers. However, the individual role-played by each event, their order in tumor dynamics and selection of downstream mutations were distinct in all three anatomical locations, with some similarities noted between LCC and rectal cancer. We found a potentially targetable alteration APC R1450* specific to RCC that has not been previously described. Differential gene expression analysis revealed multiple genes within the homeobox, G-protein coupled receptor binding and transcription regulation families were dysregulated in RCC, LCC, and rectal cancers and may have a pathological role in these cancers. Further, using a novel in silico proteomic analytic tool developed by our research group, we found distinct central or hub proteins with unique interactomes in each location. Protein expression signatures were not necessarily concordant with the tumor profiles obtained at the DNA and RNA levels, underscoring the relevance of post-transcriptional events in defining the biology of these cancers beyond molecular changes at the DNA and/or RNA level. Ultimately, not only tumor location and the respective genomic profile but also protein-protein interactions will need to be taken into account to improve treatment outcomes of colorectal cancers. Further studies that take into account the alterations found in this study may help in developing more tailored, and perhaps more effective, treatment strategies.

**Author summary:** Patients with right-sided colon cancer (RCC) has a worse prognosis compared to left-sided colon cancer (LCC). Recent data has also shown that wild-type RAS metastatic RCC’s have poor outcomes when treated with the combination of chemotherapy and anti-EGFR therapy compared to LCC and rectal cancers. Therefore, There is an urgent unmet need to understand the molecular differences between RCC, LCC, and rectal cancers. In this study, we demonstrate clonal evolutionary trajectory and the order of mutations in RCC, LCC, and rectal cancers are distinct with some similarities between LCC and rectal cancers. The order of the mutations that lead to the acquisition of crucial driver alterations may have prognostic and therapeutic implications. We also discovered a novel targetable alteration, APC R1450* to be significantly enriched in early, late and metastatic RCC but not in LCC and rectal cancers. Amazingly, proteomic signatures were discordant with DNA and RNA levels. These distinct differences in DNA, RNA and post-transcriptional events may contribute to their unique clinicopathological features.

**Conflict of Interest Statement:** **Ashiq Masood** Advisory board and speaker Bureau Bristol-Myers Squibb and Boehringer Ingelheim

**Janakiraman Subramanian** Advisory board - Astra Zeneca, Pfizer, Boehringer Ingelheim, Alexion, Paradigm, Bristol-Myers Squibb Speakers Bureau - Astra Zeneca, Boehringer Ingelheim, Lilly Research Support - Biocept and Paradigm

**Arif Hussain** Advisory board – Novartis, Bayer, Astra Zeneca Consultant – Bristol-Myers-Squibb All other authors have no conflict of interest.

## Introduction

Although often grouped as one disease, right-sided colon cancer (RCC) and left-sided colon cancer (LCC) represent clinically distinct entities. RCC is typically associated with peritoneal disease, aggressive tumors and poor prognosis.[1,2] Rectal cancer is often grouped with colon cancer. Given its proximity and continuance with the left colon, rectal cancers are commonly thought to be the same disease as colon cancers. However, the treatments for early-stage rectal cancer and colon cancer are different, in large part due to the increased rate of recurrence of rectal cancers.[3] A retrospective subgroup analysis of the landmark CALGB/SWOG 80405 trial, demonstrated significant differences in outcomes of right- and left-sided colon cancers with chemotherapy.[4] However, the study included rectal cancers within the left-sided colon cancer group in the analysis. By contrast, another subgroup analysis of a landmark trial, FIRE-3, compared colon cancers (right-sided and left-sided) to rectal cancers and found differences in treatment response between the two groups.[5] Molecular investigations of colon and rectal cancers by Sanz-Pamplona et al.[6] and Hong et al.[7] have found some differences in the mutational frequencies of the HOX, APC, ERBB2, STK11 and TP53 genes.

Using high-throughput sequencing, The Cancer Genome Atlas (TCGA) sought to molecularly characterize colorectal cancers.[8] While valuable data was obtained, this study did not actively pursue differentiating RCC from LCC and rectal cancers. Other studies that have looked at differences between RCC and LCC included the transverse colon in their definition of right-sided colon cancers.[9,10] Given that embryologically the right colon and only the first two-thirds of the transverse colon have a common origin, including the entire transverse colon as part of the right colon may lead to conflicting results. Other studies have included rectal cancers in their analysis of left-sided colon cancers.[10,11]

Given the above limitations in the published literature, in the current study we used The Cancer Genome Atlas (TCGA)[8,12] and Memorial Sloan Kettering Cancer Center (MSKCC)[11] data to evaluate differences at the DNA, RNA, microRNA and protein levels to better understand the molecular signatures that characterize RCC (cancer originating from cecum, ascending colon, hepatic flexure), LCC (cancer originating from splenic flexure, descending colon, sigmoid colon), and rectal cancers. Specifically, we sought to examine differences in clonal evolution using Pipeline for Cancer Inference (PiCnIc) on cross-sectional data.[13] Despite the many recent molecular colorectal cancer studies, to our knowledge none have specifically explored ensemble gene level clonal evolution differences and proteomic differences among RCC, LCC and rectal cancers. We employed the ConsensusDriver algorithm[14] that takes advantage of combining multiple previously described algorithms to increase driver mutation discovery. We also examined amino acid hotspots, DNA copy number, RNA, miRNA and protein expressions in these various tumors. This is the first study to systemically examine the molecular basis underlying the observed clinical differences in colorectal cancer subtypes. Our study revealed significant differences between RCC, LCC and rectal cancers that may have therapeutic relevance in these highly complex and heterogeneous tumors.

## Results

### Clonal evolution trajectories

Cancer development is a dynamic evolutionary process and the acquisition of somatic alterations often occurs in steps, with the genes interacting with each other epistatically.[15–17] After sequencing thousands of tumor samples over the last decade, it is now clear that linear progression models are in fact too simple a representation of the actual evolutionary process in tumor development.[16] Thus, understanding the mutational timing and evolutionary trajectory of tumors is key to delineating the molecular underpinnings of cancer development and progression.[15] Further, the order in which tumors acquire driver alterations can have prognostic and therapeutic implications.[18]

One strategy to study tumor evolution is by employing cross-sectional data from large patient datasets. To this end, we applied PiCnIc to our data to extract ensemble-level cancer progression models to understand the evolutionary mutational trajectories between RCC, LCC and rectal cancers.[13] All three cancer locations had mutations in APC, TP53 and KRAS, possibly reflecting common initiating somatic events (Fig 1; S1-4 Tables). Analysis revealed hierarchies of mutations in other genes that clustered around APC, TP53 and KRAS (Fig 1). In RCC, APC somatic mutations and TP53 somatic mutations were independent events. APC somatic mutations ‘selected’ for KRAS mutations or amplifications. APC somatic mutations also ‘selected’ for SMAD4 somatic mutations or deletions, BRAF mutations and amplification. KRAS and BRAF showed mutual exclusivity. Interestingly, alterations in FBWX7, TCF7L2, and SMAD2 clustered in RCC tumors harboring APC and PIK3CA mutations. With respect to TP53, alterations in this location were associated with CTNNB1, MYC or/and BRCA2 mutations.

**Fig 1.**
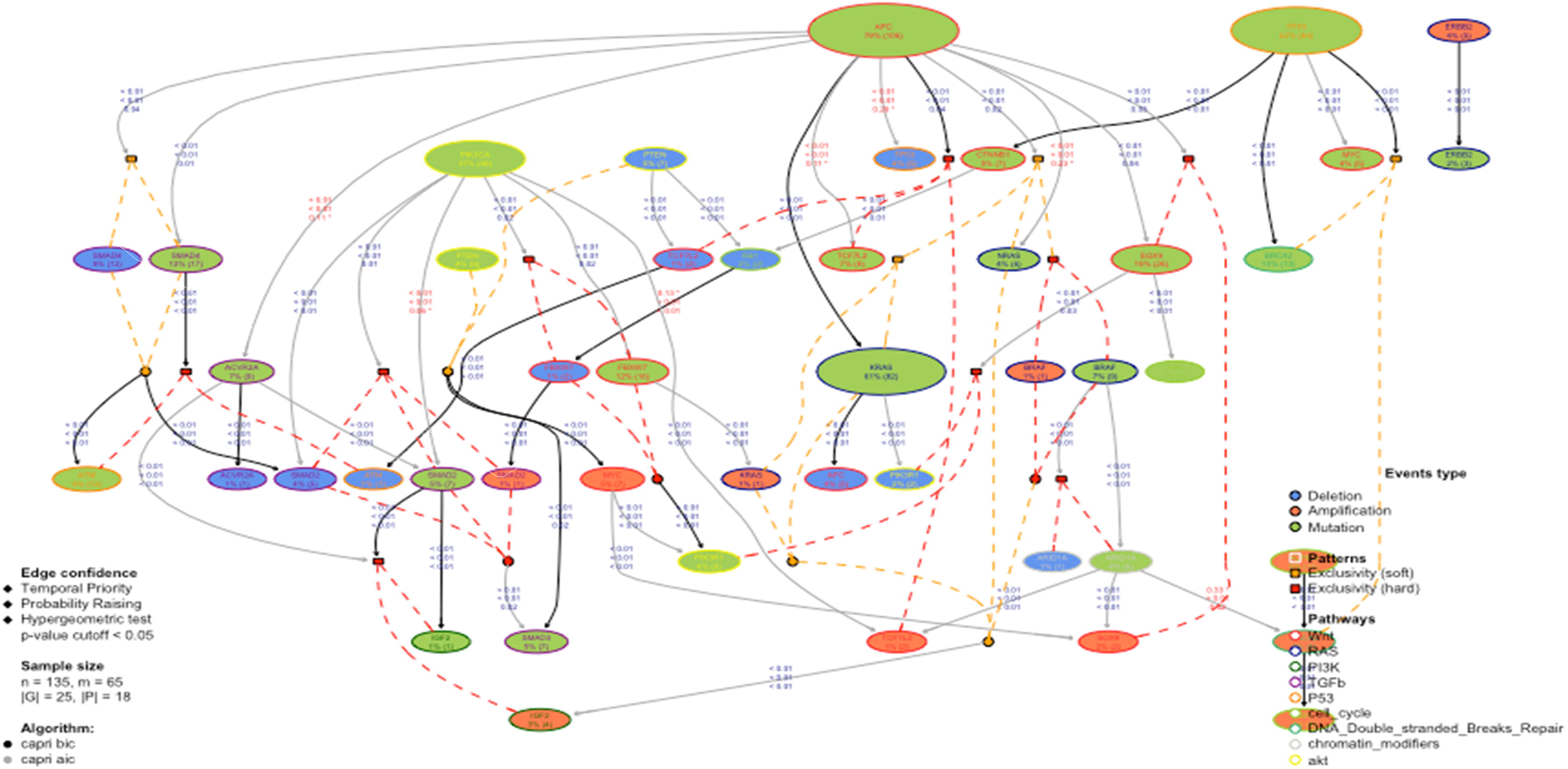
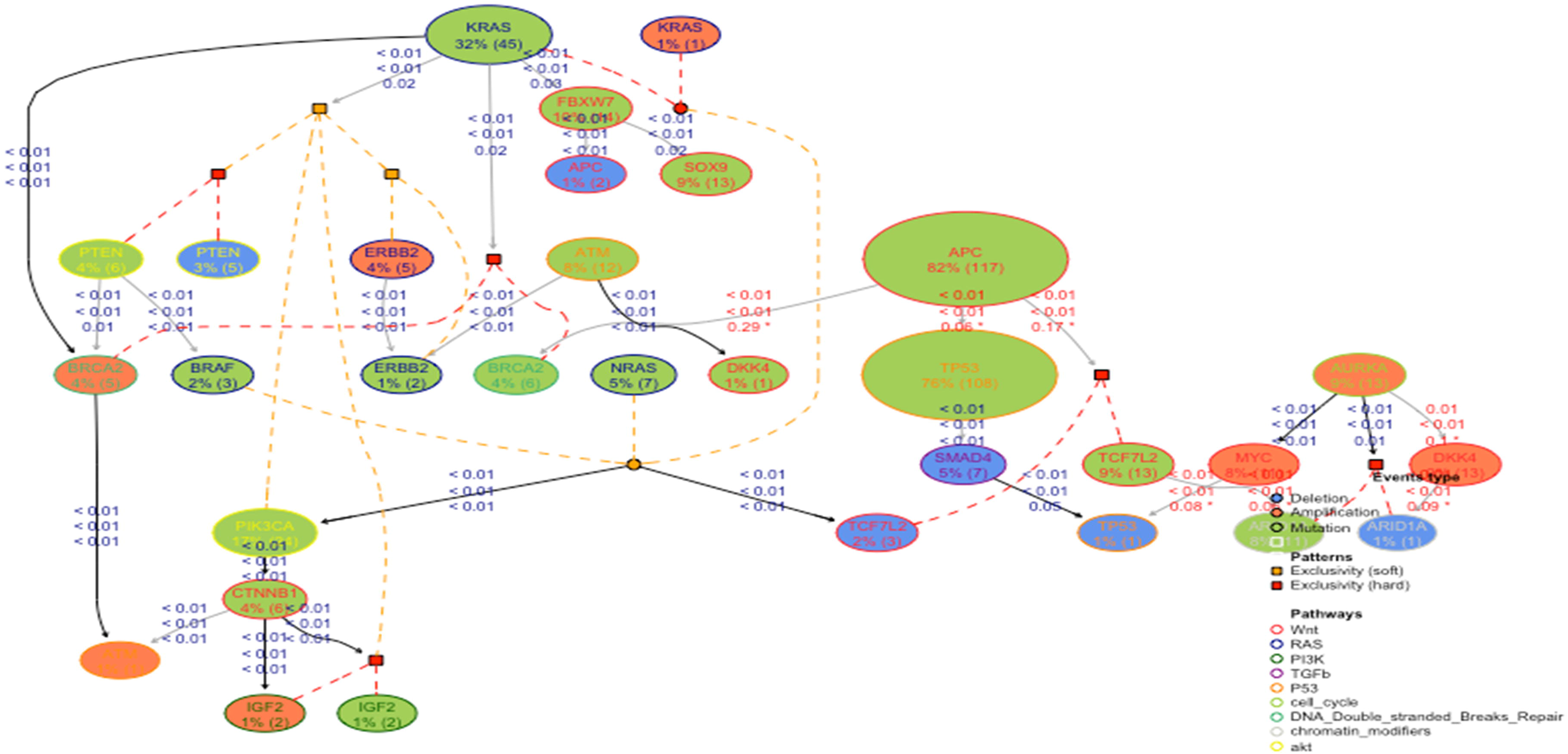
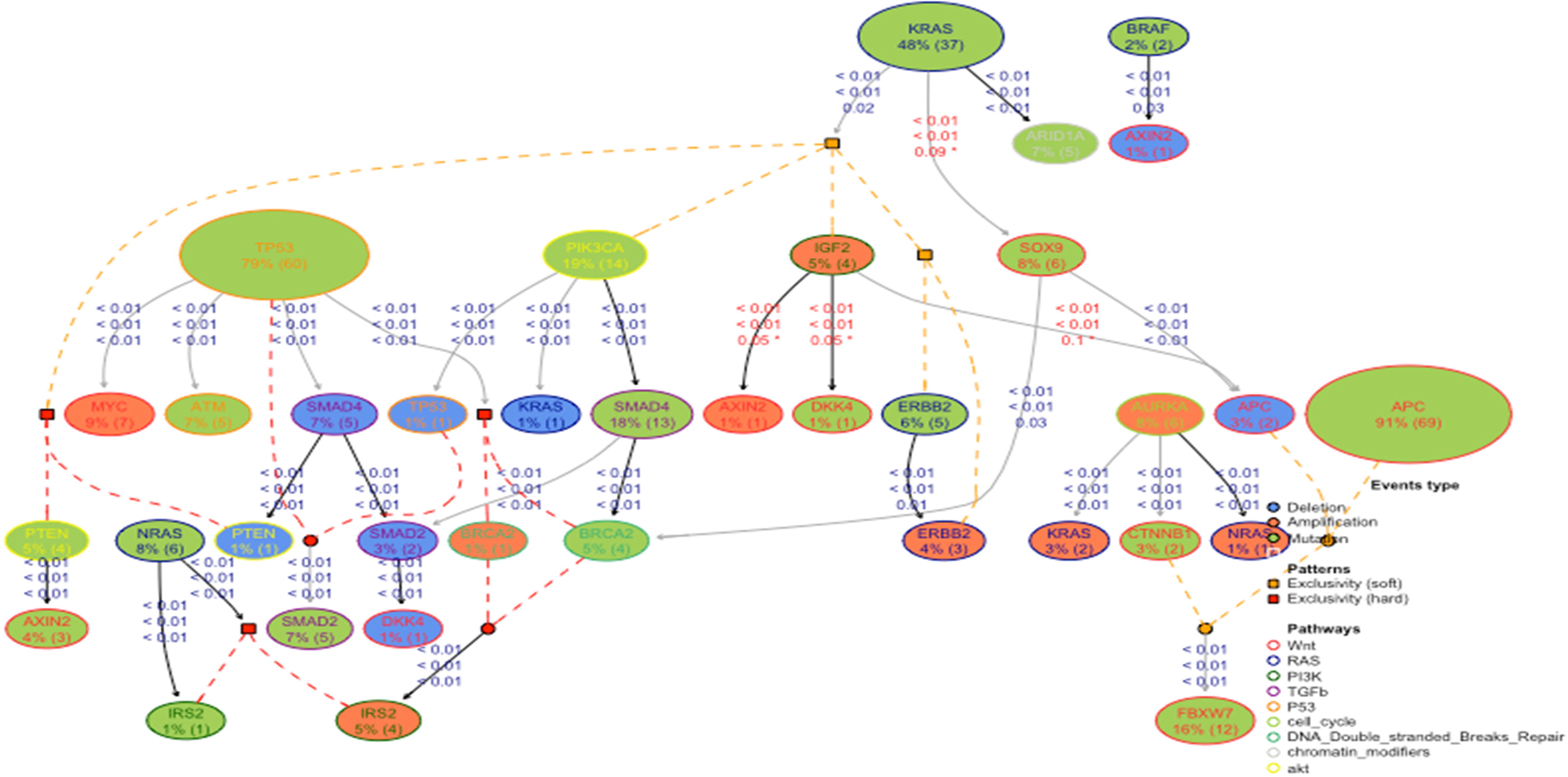
Progression model of colorectal cancers. Circular formula selects for a downstream node and logical connectedness are squares. Clonal progression events are connected with dashed lines with orange color representing soft and red color representing hard exclusivity. The p-values for temporal priority and probability raising are shown in the figure. Red p-value indicates minimum threshold above 0.05. 1a) Progression model of right-sided colon cancers, 1b) Progression model of left-sided colon cancers and 1c) Progression model of rectal cancers.

In LCC, KRAS somatic mutations ‘selected’ for BRCA2 amplification, PTEN deletions or somatic mutations, PIK3CA somatic mutations, IGF2 amplification or somatic mutations and ERBB2 amplification or somatic mutations. Unlike RCC, alterations in PIK3CA were a late event in LCC and IGF2 amplification via CTNNB1. Another striking part of the model was AURKA amplification ‘selecting’ for MYC amplification. APC seemed to ‘select’ for TP53 but this did not reach statistical significance (p = 0.06). Similarly, APC somatic mutations ‘selected’ for BRCA2 mutations (p = 0.3) and TCF7L2 somatic mutations or deletions (all p = 0.2) but this association also did not reach statistical significance (Fig 1b and S1b Fig).

In rectal cancers, key initial mutations are split between TP53 and KRAS. TP53 ‘selects’ for MYC amplification, SMAD4 deletion and BRCA2 somatic mutation or amplification. KRAS ‘selects’ for PTEN deletion or somatic mutations, PIK3CA somatic mutations, IGF2 amplification and ERBB2 amplification or somatic mutations. Among rectal cancer patients with AURKA mutations there is clustering of NRAS amplifications (Fig 1c and S1c Fig).

Our model shows significant differences in the mutational profiles of genes between RCC and LCC; the early common somatic gene mutations are associated with the ‘selection’ of different subsequent genomic events in RCC compared to LCC. The data also suggest that although LCC and rectal cancers have some similarities in the tumor progression model wherein KRAS ‘selected’ for several genes in common (such as PIK3CA, IGF2, and ERBB2 alterations), significant differences were also noted between these two sites (Figs 1b and 1c).

Taken altogether, our results show non-adherence to the established Vogelstein linear progression model of colorectal cancer progression from normal mucosa to adenoma to carcinoma. Further, the current data suggest that RCC, LCC and rectal cancers have distinct mutational behavior in the context of their evolutionary trajectories and mutational timing during cancer development and progression although initial events such as APC gatekeeper mutation appear to be similar.

### Somatic driver alteration analysis

The ConsensusDriver algorithm identified 25 significantly mutated genes (≥5% of tumors) in all three tumor locations, including mutations in the WNT, P53 and TGF-b pathways, in agreement with other studies.[8,10,11,19] In addition, we identified several novel driver mutations within the DNAH8, DST, PAPPA, and TNR genes (Fig 2).

**Fig 2.**
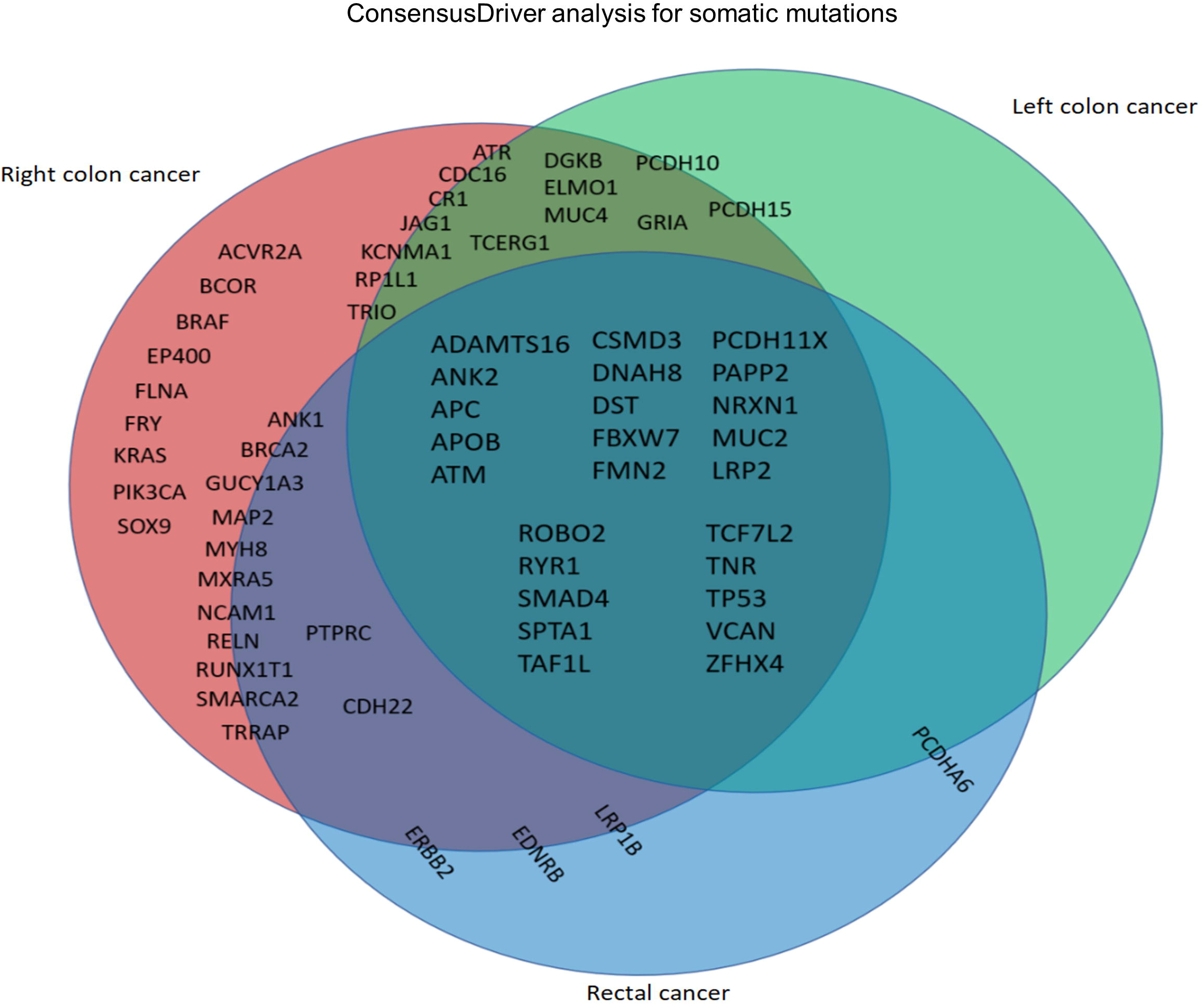
ConsensusDriver analysis for somatic mutation enrichment for left-sided, right-sided and rectal cancers. Genes located within each area met criteria for statistical significant mutation enrichment (frequency ≥ 5%, chi-square p <0.05) compared to the other two anatomical locations (e.g. KRAS mutations were found statistically enriched in right colon cancers compared to left colon cancers and rectal cancers). Genes located on the borders of the Venn diagram had enrichment for mutations in both locations but only met statistical significance for enrichment to one site but not both (e.g. ERBB2 mutations were enriched in rectal cancers compared to left colon cancers with statistical significance but not to right colon cancers. They were enriched in right colon cancers compared to left colon cancers but not to statistical significance).

Comparison of significantly mutated genes (SMGs) between RCC, LCC and rectal cancers identified 9 SMGs that were significantly enriched (≥5% of tumor samples, p<0.05) in RCC. These included RTK/RAS pathway genes: KRAS and BRAF; IGF/PI3K pathway gene: PIK3CA; WNT pathway gene: SOX9; TGFB pathway gene: ACVR2A; and MYC-pathway gene: EP400. Novel genes significantly enriched include: FRY, FLNA and BCOR. FLNA and BCOR are associated with tumor progression and metastasis[20] whereas FRY encodes a microtubule-binding protein which is conserved across species[21] (Figs 3a-c; S2 Fig; S5 and S6 Tables).

**Fig 3a.**
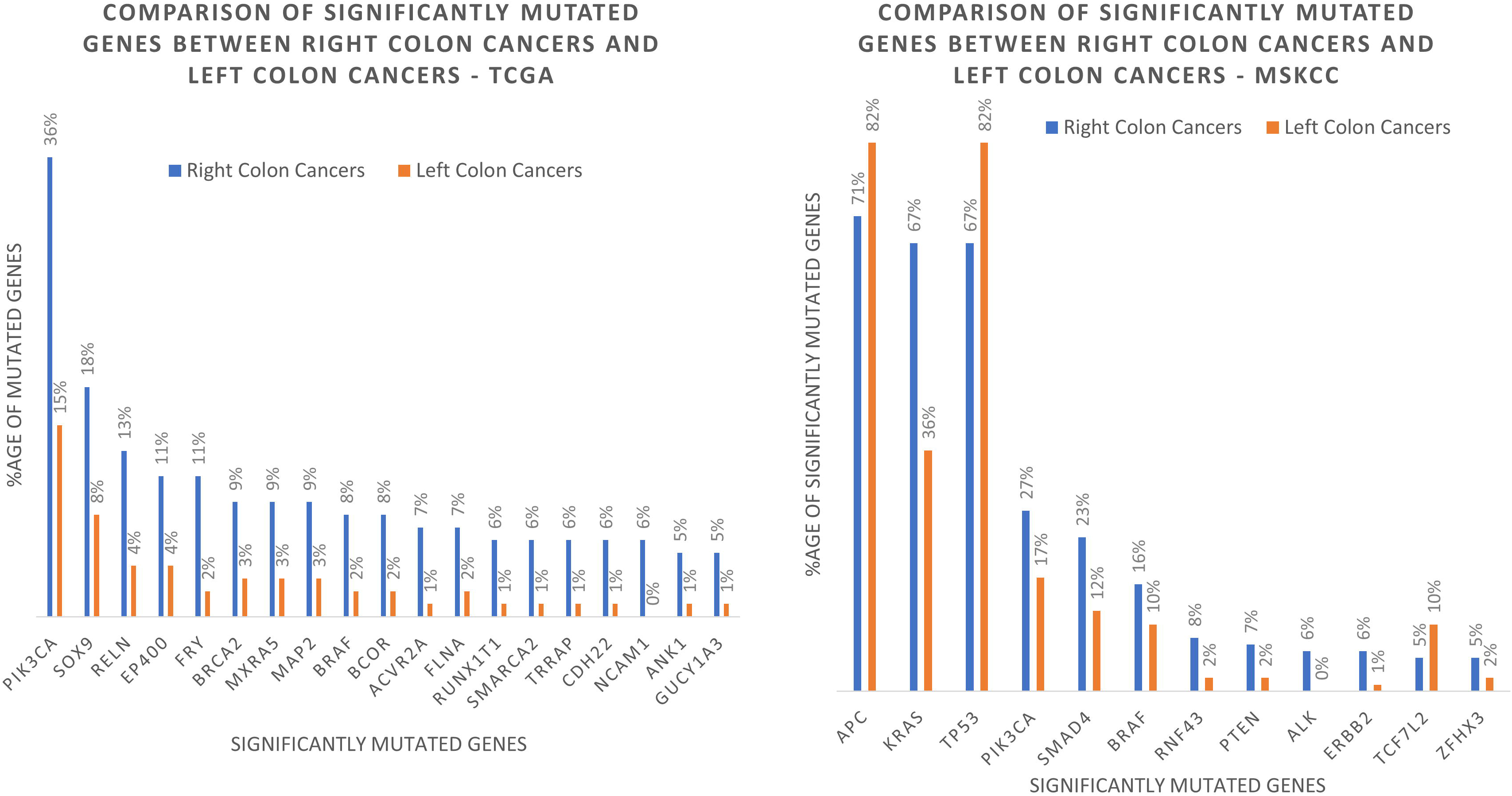
Significantly mutated genes (SMGs) between TCGA and MSKCC data set for right-sided colon cancers and left-sided colon cancers. SMGs were defined as presence of genes in ≥ 5% of tumor samples with p <0.05. Twenty SMGs were enriched in RCC compared to LCC in TCGA and twelve SMGs were enriched in MSKCC. Three SMGs (KRAS, PIK3CA, BRAF) were common between the two data sets that were enriched in RCC.

**Fig 3b.**
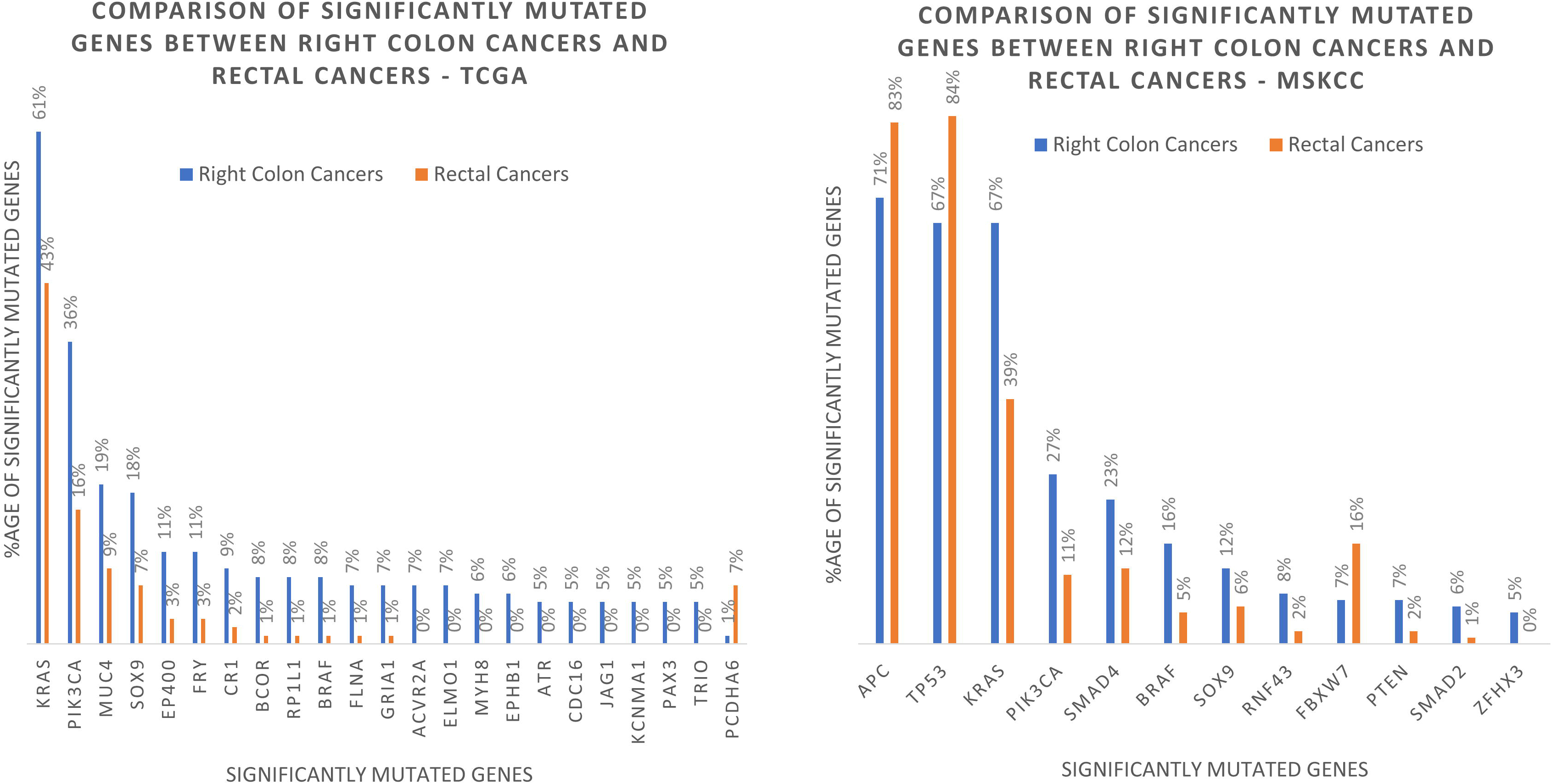
shows the SMGs between TCGA and MSKCC data set for right-sided colon cancers and rectal cancers. Twenty three SMGs were enriched in RCC vs. rectal cancers in TCGA and 12 SMGs in MSKCC. Four SMGs (KRAS, PIK3CA, SOX9, BRAF) were common between the TCGA and MSKCC data sets that were enriched in RCC.

**Fig 3c.**
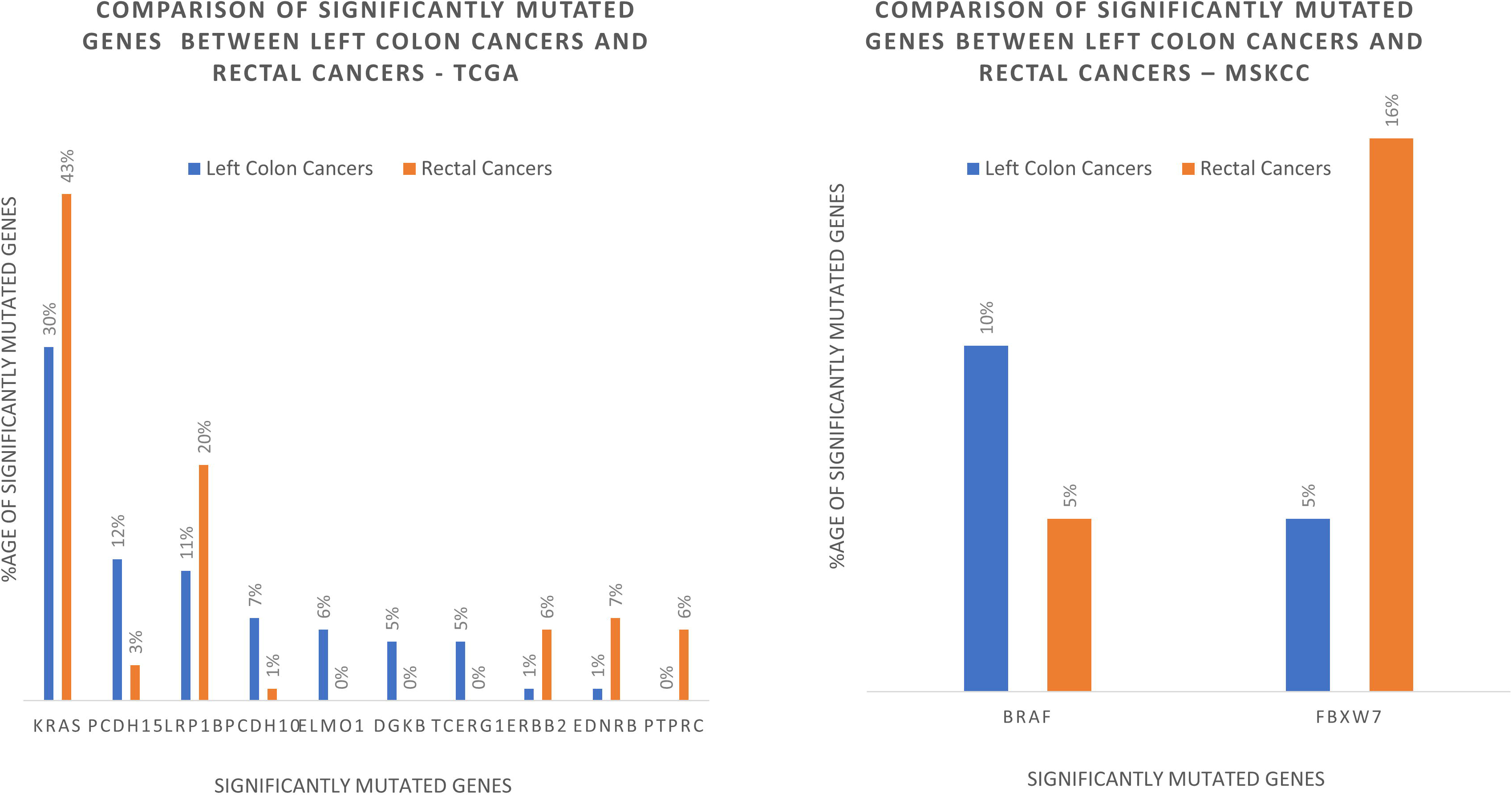
shows SMGs between the TCGA and MSKCC data sets for LCC and rectal cancers. 10 SMGs were identified in TCGA and 2 SMGs were identified in MSKCC data set. No similar SMGs were identified between the two data sets.

We also discovered somatic alterations in genes likely associated with tumor invasiveness and progression that are enriched in RCC compared to LCC (all p<0.04) but not rectal cancers (all p> 0.05): RELN,[22] MAP2,[23] NCAM1,[24] and RUNX1T1.[25] This discrepancy may be due to sample size differences (RCC n = 142, rectal cancer n = 89). Altogether, these genes are mutated in 37% of RCC (52/142) versus 10% of LCC (15/156) and showed a tendency towards mutual exclusivity (S3 Fig). TRRAP was also significantly enriched in RCC (6%) compared to LCC (1%, p < 0.02) but not rectal cancers (2%, p = 0.2). Recently, TRRAP was found to be essential for regulating p53 mutant levels in lymphomas by preventing its degradation.[26] Given its enrichment in RCC it may play a potential role to that effect in colorectal cancer.

Conversely, only one SMG was enriched in LCC, PCDH10. This gene was statistically enriched compared to rectal cancer (7% vs. 1%, p = 0.038) but was not enriched compared to RCC (7% vs. 4%, p = 0.18). This gene is a tumor suppressor gene shown to be an independent predictor of colorectal metastasis.[27]

Compared to LCC, rectal cancers were enriched for LRP1B (20% vs. 11%, p = 0.045). LRP1B functions as a LDL receptor-related protein involved in promoting growth and migration in colorectal cancers.[28] However, LRP1B did not reach statistical significance when comparing LCC to RCC (20% vs. 16%, p = 0.4335). Similarly, ERBB2 somatic mutations were significantly enriched in rectal cancers compared to LCC (6% vs. 1%, p = 0.015) but not RCC (6% vs. 2%, p = 0.15).

In addition, we also analyzed the genomic differences between RCC, LCC and rectal cancer among a cohort of patients with metastatic CRC (represented by the MSKCC cohort.[11] Mutational analyses were done with the 468 gene MSK-IMPACT clinical sequencing platform. Several genes enriched in the TCGA RCC cohort are not in the MSKCC gene panel and thus were not included in the analysis. These genes include: FRY, NCAM1, RELN, CDH22, ACVR2A, SMARCA2, TRRAP, MXRA5, EP400, FLNA, MAP2 and RUNX1T1. As in the early stage tumors from the TCGA cohort, KRAS and PIK3CA mutations were enriched in metastatic RCC compared to LCC and rectal cancers (all p <0.005) suggesting that they may have a larger role to play in RCC development. Similarly, BRAF was highly enriched in RCC compared to LCC and rectal cancers in both data sets (all p <0.04). Interestingly, ZFHX3, mutated at low rates in the TCGA cohort was significantly enriched in metastatic RCC (5%) compared to metastatic LCC (2%) and metastatic rectal cancers (0%) (all p <0.03) (Figs 3a-c).

TP53 was more frequently mutated in LCC (82%) and rectal cancers (84%) than RCC (67%) in the MSKCC cohort (both p <0.001). This trend was also seen in the TCGA cohort, though not to statistical significance (71% LCC vs. 61% RCC, p = 0.07; 71% rectal cancer vs. 61% RCC, p = 0.14). LCC was also highly enriched for APC mutations compared to RCC in the advanced stage CRC cohort (82% vs. 71%, p = 0.002), whereas APC mutations were equally distributed between RCC and LCC in the TCGA data set (78% vs. 77%, respectively). FBXW7 mutations were enriched in rectal cancers (16%) compared to either LCC or RCC (5% and 7% respectively, both p <0.01) (Figs 3a-c).

Next, we analyzed the concordance or discordance of driver mutations between biopsies taken from primary tumor sites vs. metastatic sites within the MSKCC CRC cohort. We found minimal differences in driver mutations between the primary sites and metastatic sites (S7 Table). Only 5 genes, mostly among RCC made our statistical cut off for enrichment (mutation frequency ≥ 5%, p <0.05). In RCC, primary tumor biopsies were enriched in ERBB2 and ARID1A mutations (11% vs. 1%, p = 0.0021; 8% vs. 1%, p = 0.0163, respectively), whereas metastatic site biopsies had higher enrichment for NRAS and EPHA5 mutations (6% vs. 0%, p = 0.02; 9% vs 2% p = 0.02, respectively; S4a Fig). In rectal cancers, FBXW7 mutations were enriched in primary site biopsies compared to metastatic sites (5% vs 0%, p = 0.03; S4b Fig). Surprisingly, no differences were seen in driver mutation enrichment in LCC when comparing primary site biopsies to metastatic site biopsies (S7 Table).

Although not identified as a significantly mutated gene by ConsensusDriver, we analyzed the distribution of AMER1 mutations in colorectal cancers as recent studies have identified such mutations in CRC.[8,29] We found this gene to be significantly enriched in RCC compared to LCC and rectal cancer in the TCGA data set (both 23% vs. 3%, p <0.0001). Mutations (mostly truncating) in AMER1 and other keys genes of the β-catenin destruction complex were mutually exclusive of each other (S5a-c Figs).

However, these findings were not replicated in the MSKCC data set.

### Mutation hotspot analysis

We studied somatic mutations at the residue sites that can disrupt functional protein domains leading to tumorigenesis and clonal evolution via selective pressure.[30] As an example, KRAS G12 and G13 are commonly known hotspots in multiple cancers including colorectal cancer. Mutations at these residues affect the protein’s ability to hydrolyze GTP leading to constitutive activity and downstream signaling.[31] We found these mutations were enriched in RCC compared to LCC and rectal cancers in both the TCGA and the metastatic MSKCC data (all p <0.05). In addition, we found APC R1450* to be a significant mutation in colorectal cancers. Specifically, APC R1450* was significantly enriched in RCC (TCGA 15%, MSKCC 12%) compared to LCC (TCGA and MSKCC 1%) and rectal tumors (TCGA and MSKCC 1%) in both the TCGA and MSKCC datasets (all p <0.001, Figure 4). To our knowledge, this is the first report to describe the APC R1450* mutation as being predominantly located in RCC. This particular hotspot in APC is exclusively a truncation mutation and lies within the MCR domain of the protein, which is a highly mutated area. The resulting truncated mutant conserves beta-catenin binding sites (15 AA repeats) but loses all three axin-binding sites (SAMP repeats) and microtubule interaction via EB1 and PdZ domains [32]. The MCR domain was previously defined as codons 1282 – 1581.[33,34] Unlike APC R1450*, the frequency of other mutations within this region is relatively similar among the TCGA and MSKCC data sets. The relative frequencies of non-R1450* mutations within the MCR domain of APC for RCC were 63% and 64% in the TCGA and MSKCC data sets, respectively, for LCC 52% and 51%, respectively, and for rectal cancers 64% vs 58% (which did not meet statistical significance, p=0.35). Another gene of interest is SMAD4. Specific to the TCGA samples, SMAD4 D537 missense mutations were highly enriched in rectal cancers compared to LCC and RCC (both p <0.03). Previously seen in 3% of colorectal cancers,[35] we found SMAD4 D537 to present almost exclusively in rectal cancer (rectal cancers 6%, LCC 0%, RCC 1%) in the TCGA dataset. Mutations in BRAF V600 were significantly enriched in colon cancers (RCC 11%, LCC 6%) compared to rectal cancers (1%) in the MSKCC data set (both p <0.007). Taken together, these emerging data suggest that certain mutations are enriched for in the TCGA patient samples but not the MSKCC samples, or vice versa, while other mutations may be common to both. The TCGA samples are mainly from patients who had earlier stage CRC, whereas the MSKCC samples represent primary tumor sites or metastatic biopsies from patients who also had metastatic disease. Whether the different distribution patterns of some of the genetic changes noted in the TCGA and the MSKCC patient populations contribute in part to the differences in their underlying clinical features is an intriguing question.

**Fig 4.**
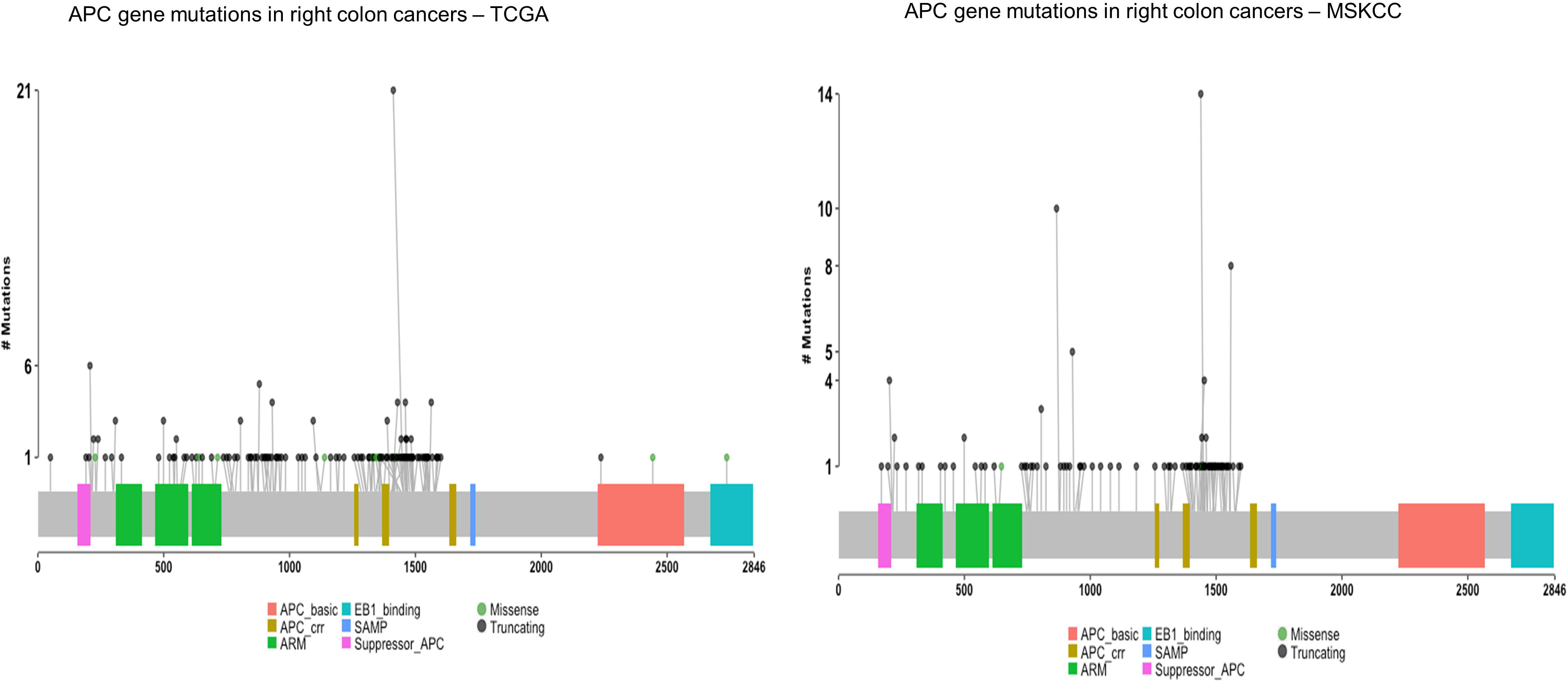
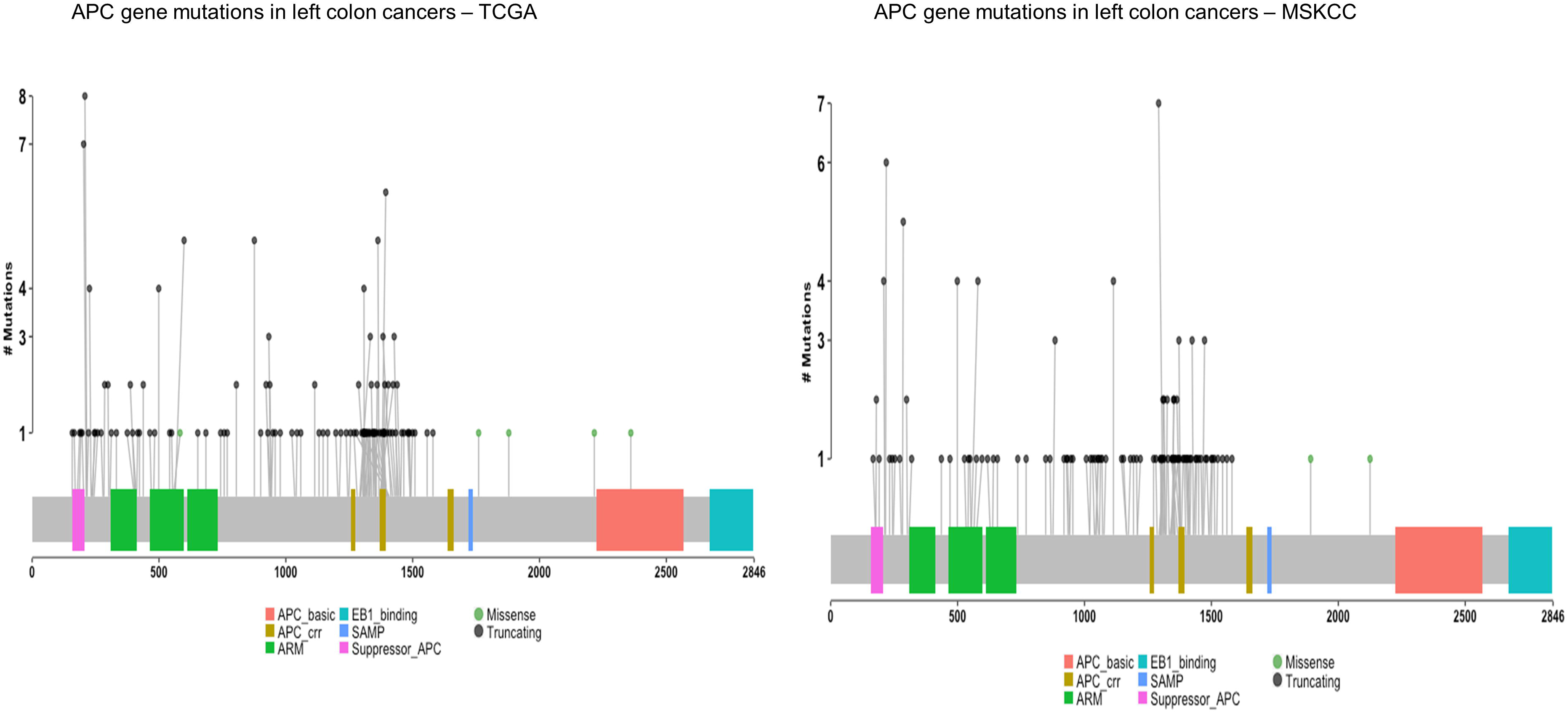
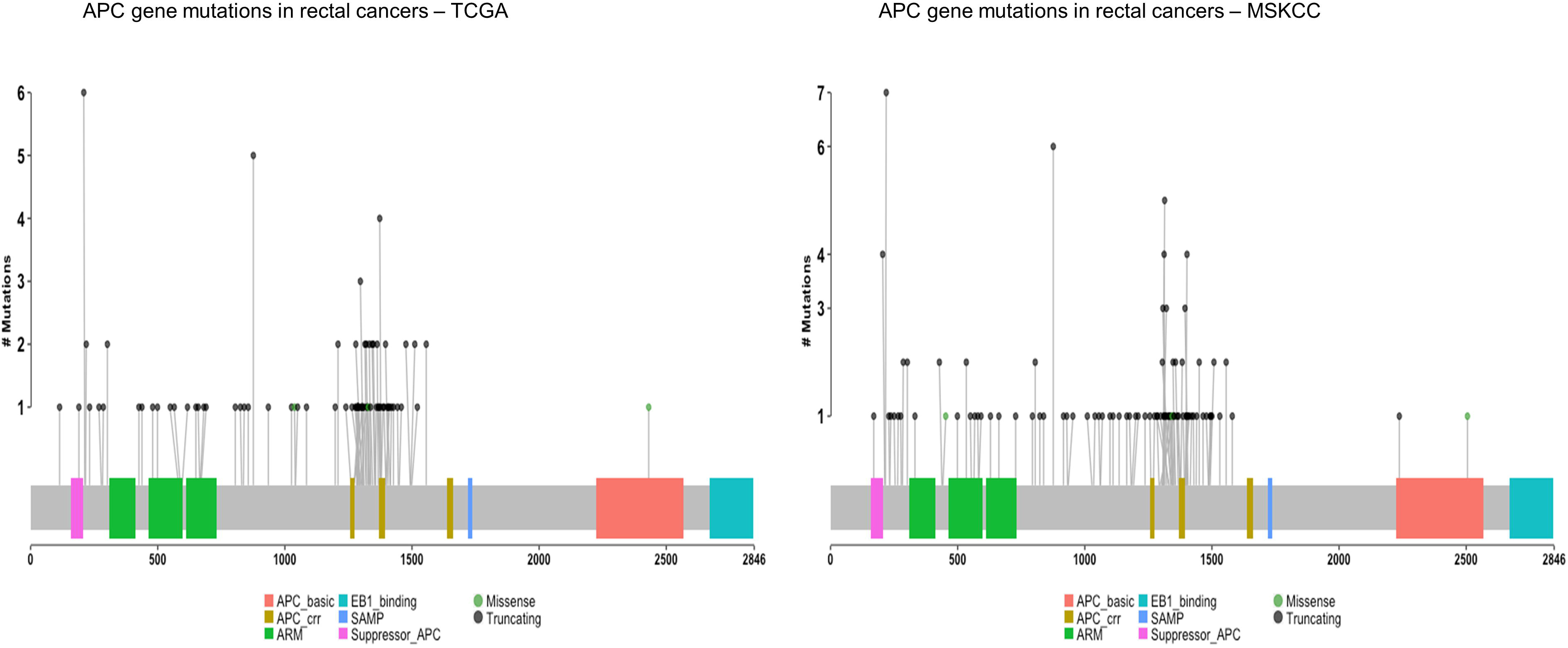
shows the frequency of the APC hotspot R1450 residue in (a) right-sided colon cancers, (b) left-sided colon cancers and (c) rectal cancers in TCGA (left) and MSKCC (right) datasets. Y-axis represent total number of mutations at each residue.

### Differences in somatic copy number alterations (SCNAs) between RCC and LCC

Our somatic copy number analysis identified 13 unique amplified regions in LCC compared to RCC. Conversely, only 2 unique regions were amplified in RCC. The RCC amplified regions correspond to ZNF217 (a zinc finger protein). The LCC amplified regions correspond to KLF5 (zinc finger transcription factor). In addition, 13 unique deleted regions were identified in LCC compared to RCC where only 4 regions were deleted (S6a and S6b Figs). The deleted regions in LCC correspond to 6 known oncogenes or tumor suppressor genes: SMAD3, STK11, CASP3, BRCA1, NF1 and MAP2K4. No candidate genes were identified in the deleted regions in RCC (S8 Table).

### Differences in RNA expression between RCC, LCC and rectal cancers

RNA expression analysis from TCGA revealed 124 differentially expressed genes (DEG) between RCC and LCC (p-value <0.05, log2 ratio > 1.5); among these 82 were upregulated and 42 downregulated (S9a and S9b Tables). Similarly, 146 genes were differentially expressed between RCC and rectal cancers (p <0.05, log2 ratio > 1.5); among these 84 genes were upregulated and 62 downregulated (S10a and S10b Tables). Despite the close anatomic proximity and similar embryological origin of LCC and rectal cancers, several differences at the gene expression level were also noted between the two. In particular, 38 genes were differentially expressed between LCC and rectal cancer, including 22 genes that were upregulated and 16 that were downregulated (S11a and S11b Tables). 69 differentially expressed genes were shared by all three anatomical locations, including 44 upregulated and 25 downregulated genes (Fig 5).

**Fig 5.**
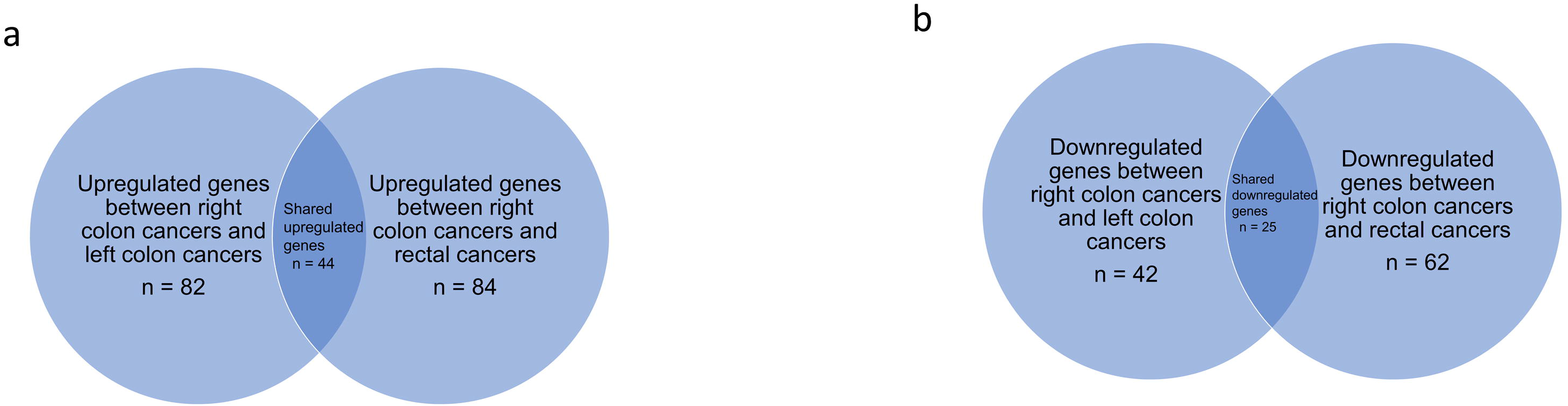
shows upregulated and downregulated genes between RCC, LCC and rectal cancers. a) upregulated genes and 44 upregulated genes shared among the three anatomical sites. b) downregulated genes and 25 downregulated genes shared among the three anatomical sites.

In an effort to better understand the potential relevance of the above changes in gene expression, we performed Functional Annotation Clustering using DAVID[36,37]. The results are shown in S12-14 Tables. The majority of clusters in our enrichment analysis were similar when comparing RCC vs. LCC and RCC vs. rectal cancer. The downregulated clusters included homeobox-related, G-protein coupled receptor binding (hormonal regulation)-related, and transcription regulation-related genes, although for the latter statistical significance was not reached between RCC and rectal cancer. Similarly, upregulated clusters in both groups showed enrichment in homeobox-related, lipid and cholesterol metabolism-related, and transcription regulation-related genes. In addition, we identified other differentially expressed gene clusters unique to RCC that were related to pathways for oxygen transport or sodium transport, or to oxidoreductase activity.

When we applied a hierarchical (agglomerative) clustering method, REVIGO[38] to our Gene Ontology (GO) biological process obtained from DAVID analysis, the RCC vs. LCC and RCC vs. rectal cancer showed dysregulation of similar pathways such as negative regulation of inflammatory response, lipid and cholesterol metabolism, keratinocyte differentiation, cell cycle - cell division control protein 42 regulation. In addition, RCC vs. LCC showed enrichment in cell differentiation and embryonic limb morphogenesis; whereas, RCC vs. rectal cancer was enriched in ion transport, sodium ion transport and somatic stem cell population maintenance among others. The LCC and rectal cancers showed no significantly altered biological process except iron transport and melanin biosynthesis (S7 and S8a-c Figs and S15-17 Tables). The novel graph theoretic clustering algorithm, Molecular Complex Detection (MCODE),[39] on protein-protein interaction network obtained from string database was performed using our gene list. RCC vs LCC Protein-Protein Interaction network showed enrichment in peptide hormone pathway, lipid metabolism and keratinization pathway (Fig 6a). Similarly, RCC vs. rectal cancers showed peptide hormonal and lipid metabolism dysregulation along with the tight junctions (Fig 6b). LCC vs. rectal cancers showed no pathway enrichment.

**Fig 6a.**
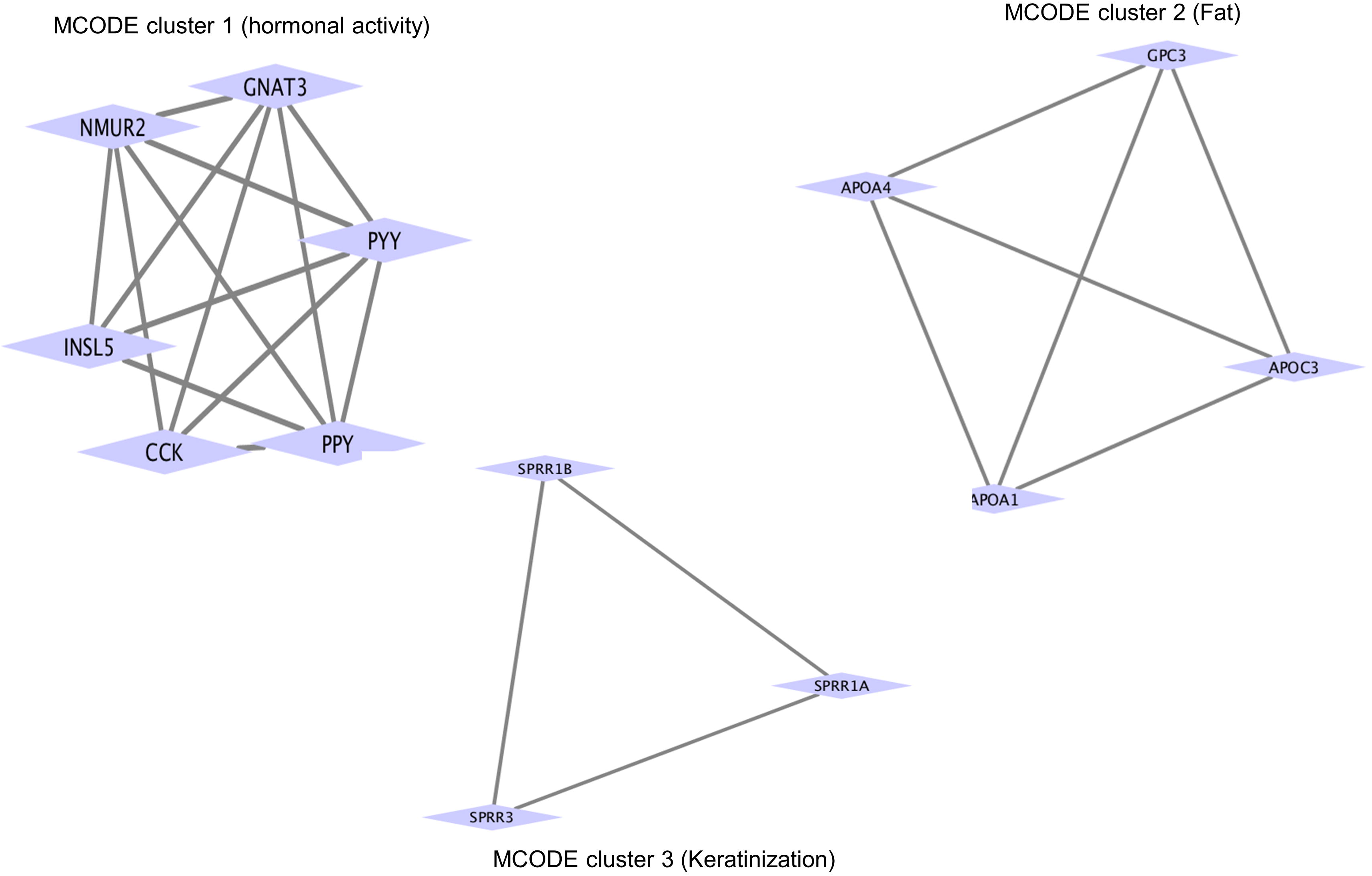
shows different MCODE clusters for upregulated and downregulated genes when right-sided colon cancers were compared with left-sided colon cancers. MCODE cluster 1 involved in hormonal activity, MCODE cluster 2 involved in fat metabolism and MCODE cluster 3 involved in keratinization.

**Fig 6b.**
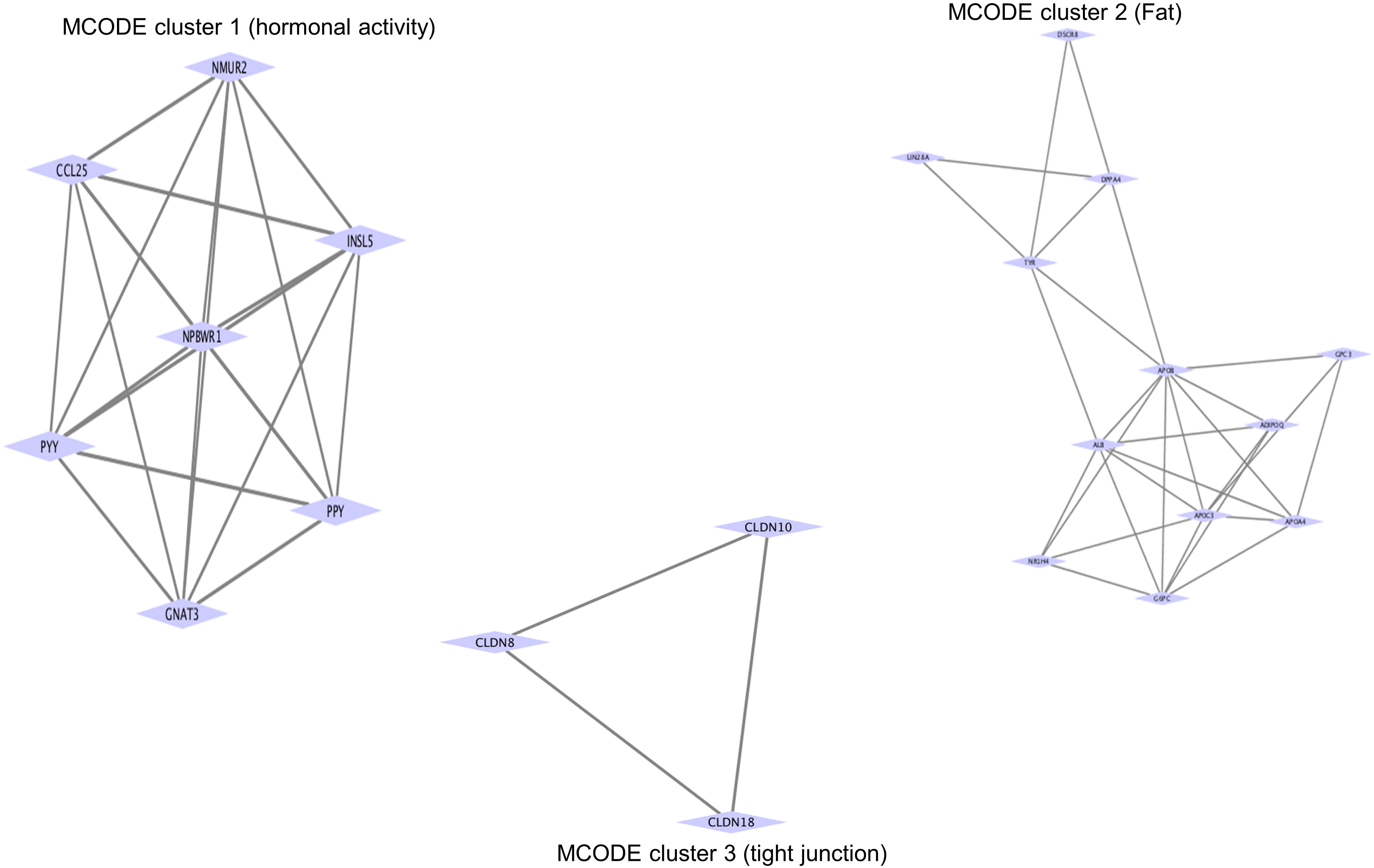
shows different MCODE clusters for upregulated and downregulated genes when right-sided colon cancers were compared with rectal cancers. MCODE cluster 1 involved in hormonal activity, MCODE cluster 2 involved in fat metabolism and MCODE cluster 3 associated with tight junction.

### MicroRNA (miRNA) expression differences between RCC, LCC and rectal cancers

We identified 13 cancer-related miRNAs that were differentially expressed (adjusted p <0.05 and fold change > 1.5) between RCC and LCC, and 31 that were differentially expressed between RCC and rectal cancers. In addition, 16 miRNAs were differentially expressed between LCC and rectal cancers (S18 Table).

Seven miRNAs (4 upregulated and 3 downregulated) were dysregulated when comparing RCC to both LCC and rectal cancer (S9a Fig, S19 Table), while 3 were upregulated and 3 downregulated when RCC was compared to only LCC (S9b Fig, S20 Table), and 11 were upregulated and 13 downregulated when RCC was compared to only rectal cancer (S9c Fig, S21 Table). The various miRNAs identified have been implicated in a range of pathological processes associated with oncogenesis (S19-21 Tables). Although further investigation of the differentially expressed miRNAs is needed to better understand and determine to what extent they may contribute to the tumor biology of RCC, LCC and rectal cancers, conceivably there may be potential for these miRNAs to serve as tumor anatomic location specific biomarkers.

### Proteomic analysis of right-sided, left-sided colon and rectal cancers

Using The Cancer Proteome Atlas (TCPA) data,[40] we examined RCC, LCC and rectal cancers by proteomic cancer co-expression subnetworks using association estimators methodology previously described.[41] Interestingly, no common protein emerged as having a centralized role (hub protein) across all 3 cancer locations. Within protein-protein interaction networks, several hub proteins, and their respective interactomes, were found to be unique to each of the locations (Fig 7).

**Fig 7a.**
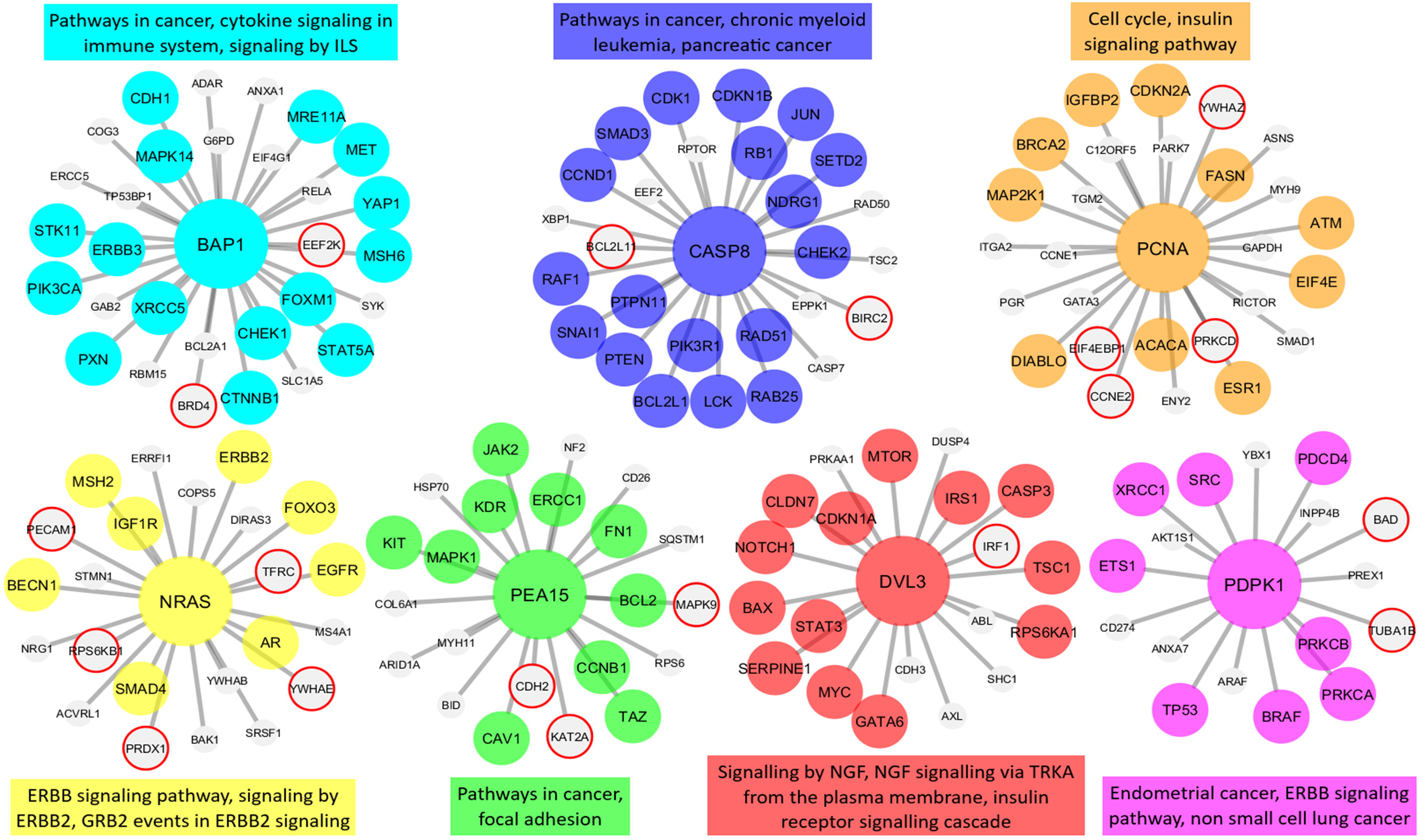
The hub genes and neighbors in the disease-related sub-networks obtained by the most successful KDE method (in terms of precision score) on COAD-RIGHT group. The genes registered in DisGeNET and experimentally confirmed for the diseases are shown with colored and larger nodes. Among these, genes that are not colored but have a red frame have a PMID value of one. There is no entry in DisGeNET for the grey colored nodes. Also, the most associated top three biological pathways, to which each module is related, are given above or below the relevant module to annotate each module.

**Fig 7b.**
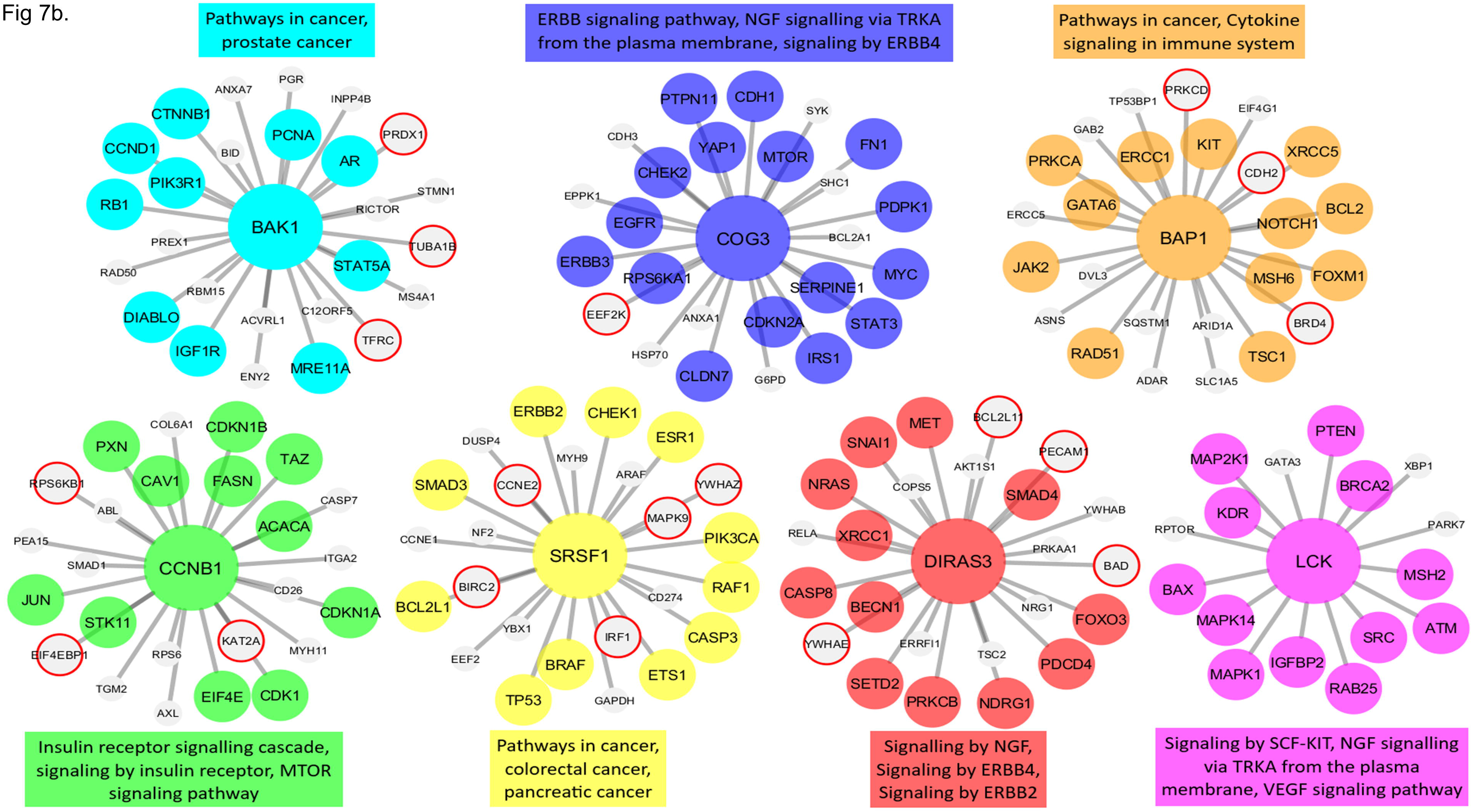
The hub genes and neighbors in the disease-related sub-networks obtained by the most successful MM method (in terms of precision score) on COAD-LEFT group. The genes registered in DisGeNET and experimentally confirmed for the diseases are shown with colored and larger nodes. Among these, genes that are not colored but have a red frame have a PMID value of one. There is no entry in DisGeNET for the grey colored nodes. Also, the most associated top three biological pathways, to which each module is related, are given above or below the relevant module to annotate each module.

**Fig 7c.**
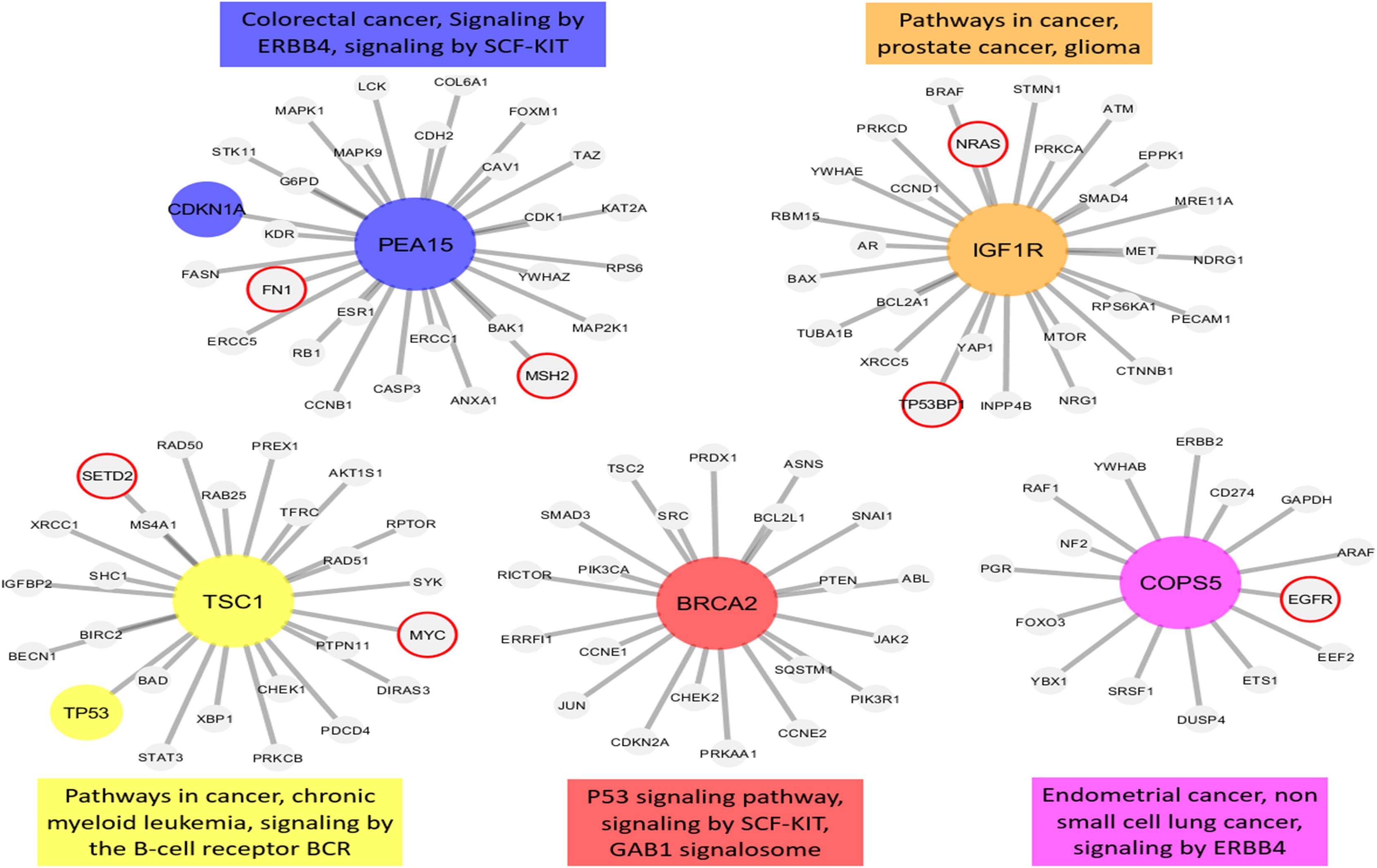
The hub genes and neighbors in the disease-related sub-networks obtained by the most successful MM method (in terms of precision score) on READ-RECTUM group. The genes registered in DisGeNET and experimentally confirmed for the diseases are shown with colored and larger nodes. Among these, genes that are not colored but have a red frame have a PMID value of one. There is no entry in DisGeNET for the grey colored nodes. Also, the most associated top three biological pathways, to which each module is related, are given above or below the relevant module to annotate each module.

As shown in Fig 7a, several hub proteins that might have a major role in RCC were identified and include: BAP1 (tumor suppressor gene) CASP8 (apoptosis) PCNA (DNA repair) NRAS (RTK-RAS pathway) PEA15 (apoptosis and RET signaling) DVL3 (cell proliferation and ATM-dependent DNA damage response) and PDPK1 (growth regulation) Among the potentially significant hub proteins in LCC were the following: BAP1, BAK1 (apoptosis and prognostic in breast cancer,[42]) COG3 (protein glycosylation/golgi function) CCNB1 (mitosis and prognosis in breast cancer) SRSF1 (RNA splicing and prognosis in small cell lung cancer) DIRAS3 (tumor suppressor gene) and LCK (resistance to apoptosis) (Fig 7b). Hub proteins unique to rectal cancers were: IGF1R (proliferation, invasion, migration), TSC1 (cell growth) BRCA2 (DNA repair) and COPS5 (multiple pathways) (Fig 7c). BAP1 was found to have a prominent role in both RCC and LCC. BAP1 is a BRCA1 associated protein that acts as a nuclear-localized deubiquitinating enzyme.[43] In vitro studies have shown that decreased protein and mRNA levels of BAP1 may be related to colorectal cancer.[43] Interestingly, although there are several conserved interactions, overall the BAP1 interactome of LCC diverges from that of RCC. Among the conserved interacting proteins are: BRD4, ADAR, GAB2, SLC1A5, EIF4G1, ERCC5 and TP53BP1, BRD4, ADAR, MSH6, FOXM1 and XRCC5. Specific to LCC, BAP1 showed interactions with ERCC1, PRKCA, GATA6, JAK2, RAD51, TSC1, RSC1, NOTCH1, BCL2, KIT, PRKCD, CDH2, ARID1A, ASNS, SQSTM1 and DVL3. Specific to RCC, BAP1 was noted to interact with CDH1, MAPK14, MRE11A, MET, YAP1, STK11, ERBB3, PIK3CA, PXN, CHEK1, CTNNB1, STAT5A, EEF2K, G6PD, COG3, RBM15, BCL2A1, SYK, RELA and ANXA1.

Altogether, our results suggest BAP1 may have an essential role in carcinogenesis of colon cancer with conserved as well as divergent evolutionary interactions with other proteins in RCC and LCC that are largely absent in rectal cancers. The hub protein PEA15 may also play an essential role in RCC and rectal cancer. It is a ubiquitous cytoplasmic protein involved in multiple cellular processes through its role in intracellular signaling.[44] The interactomes have common associations with several proteins in RCC and rectal cancer: CAV1, CCNB1, CDH2, COL6A1, ERCC1, FN1, KAT2A, KDR, MAP2K1, MAP2K9, RPS6 and TAZ. Unique to RCC, the PEA15 interactome proteins include: ARID1A, BCL2, BID, CD26, HSP70, JAK2, KIT, MYH11, NF2 and SQSTM1, while rectal cancer PEA15 interactome proteins include: ANXA1, BAK1, CASP3, CDK1, CDKN1A, ERCC5, ESR1, FASN, FOXM1, G6PD, LCK, MAP2K1, MSH2, RB1, STK11 and YWHAZ. By contrast, no common hub proteins were found between LCC and rectal cancer.

## Discussion

Anatomic location of the primary tumor plays a significant role in the choice of treatment and outcomes for patients with CRC. Earlier studies have shown significant genomic differences between RCC and LCC. Using data available from the TCGA and the MSKCC gene banks and a series of data processing algorithms, we carried out a systematic bioinformatics analysis of RCC, LCC and rectal cancers at the DNA, RNA, miRNA and protein level in an effort to identify some of the underlying molecular processes that may contribute to the respective clinical behavior of these cancers. To our knowledge, this is among the first efforts to carry out bioinformatics analysis of anatomically distinct colorectal cancers at the level of DNA, RNA and protein within the same study. In addition to using the PiCnIc algorithm to identify clonal gene associations, we carried out analyses to identify somatic driver mutations, somatic copy number changes, mutation hotspots, differential RNA and miRNA expression, and protein-protein interaction networks.

We show that despite sharing the same critical initial events involving APC, KRAS and TP53 genes, there were differences in the order, selection and significance of these key genomic events in the development of RCC and LCC. We were also able to identify significant differences in the role played by other downstream somatic alterations in right-sided versus left-sided versus rectal tumors. We show that cancer development in CRC is a highly complex process involving somatic mutational events associated with the interplay and cross-talk between critical oncogenic pathways. Further, our model shows that the differences between RCC, LCC and rectal cancers are not just due to unique genomic events but rather are also due to differences in how the key (common) initial genomic events interact and select subsequent downstream events. These unique developmental trajectories may contribute to some of the differences between RCC, LCC and rectal cancers with respect to their underlying tumor biology. Further understanding of these mutational events and their temporal and spatial relationships in the cancer evolutionary process may determine their clinical behavior, including patterns of metastasis, response to treatment and clinical outcome.

To our knowledge, this is the first study to identify APC R1450* and AMER1 mutations as significantly enriched in RCC compared to LCC and RC. We show that they are mutually exclusive from other β-catenin destruction complex genes (S5a-c Figs), suggesting that they may be early events in right-sided colon cancer tumorigenesis. The APC R1450* and neighboring truncations or frameshift mutations in the MCR region comprise the vast majority of APC mutations. Given the recent findings by Zhang et al. demonstrating the efficacy of TASIN-1(small molecule inhibitor) in a murine xenograft model of human colorectal cancer harboring a truncation mutation (A1309*) similar to APC R1450* suggests that this mutation may be biologically relevant.[45,46]

Our study uncovered several mutated genes that were enriched in RCC compared to either LCC or rectal cancers (frequency >= 5%, p <0.05). Among these we identified a novel gene not previously described in colorectal cancer, FRY. This gene encodes a microtubule binding protein that plays a crucial role in the structural integrity of mitotic centrosomes. It may play a role in the interactions of AURKA which we found in our ensemble-level progression model to be an essential event in tumorigenesis. Several other genes associated with progression and aggressiveness were enriched in RCC compared to LCC, including FLNA which has been recently implicated in CRC,[20] RELN, a regulator of neuronal migration which has previously been shown to increase invasiveness when knocked down in pancreatic cancer cell lines,[22] MAP2 which is associated with cellular adhesion and mitotic spindle assembly and shown to play a role in melanoma progression[23] and NCAM1, a neuronal cell adhesion molecule implicated in metastasis in cancers of the pancreas.[24]

We identified significant differences in the mutational profile of early stage and metastatic CRC. Whereas the TCGA dataset was produced via whole exome sequencing, the MSKCC dataset was produced via a targeted sequencing platform and thus had a defined number of genes, which limited our ability to detect novel gene mutations in the metastatic CRC cohort. Comparative analysis of RCC stage IV primary tumor and metastasis biopsy samples identified ERBB2 and ARIDA 1 enrichment in primary tumor biopsies, whereas NRAS and EPHA5 were enriched in metastatic biopsies. Discrepancies between primary and metastatic sites with respect to ERBB2 is well documented in breast cancer.[47] Similarly, understanding discrepancies in RAS mutations between primary and metastatic sites is relevant since studies have shown that CRC with wild-type RAS responds better to anti-EGFR therapy plus systemic chemotherapy compared to CRC with mutant RAS.[31]

Significant differences between RCC and LCC and RCC and rectal cancers were also observed on gene and miRNA expression analysis. Although one of the limitations of the present study was the lack of normal control samples, three different algorithms (DAVID, RAVIGO, MCODE) were in agreement and revealed significant dysregulation at the RNA level between these anatomically located tumors. Genes involved in cell growth and differentiation (HOX genes), lipid metabolism and G-protein receptor coupled hormonal signaling, among others, were differentially expressed, with the differences being particularly notable between the right-sided and left-sided cancers, whereas gene expression patterns were more congruent between LCC and rectal cancers.

For example, HOX genes have a crucial role in cell growth and differentiation during embryological development. Their role has previously been described in multiple solid tumors including RCC.[48] Another key pathway that was enriched between RCC compared to LCC and rectal cancers was lipid metabolism. Metabolic reprogramming is evolving as a hallmark of cancer, and lipid metabolism dysregulation can lead to tumor growth and metastasis.[49] Similarly, G-coupled receptor-related hormonal dysregulation is currently gaining momentum as they are believed to be related to cancer progression.[50] Multiple other studies have shown the role of cell cycle in cancer pathogenesis, and cell cycle regulators are currently being used in the clinic.[51,52]

A unique aspect of this study is that it also evaluated protein expression patterns in RCC, LCC and rectal cancers using the TCPA dataset. We identified dysregulation in several key hub proteins, their interactomes and known pathways that have been implicated in oncogenesis (Fig 7). A somewhat surprising observation from this analysis is that the protein hubs and their interactomes are distinct for each of the anatomically defined tumor sites examined. Further, these protein signatures are not necessarily concordant with the tumor profiles obtained at the DNA and RNA levels. Thus, although identifying altered genes and altered gene expressions have been of paramount importance, clarifying post-transcriptional events and protein-protein interactions will also be highly relevant to understanding the variations in tumor biology and clinical behavior of tumors. In conclusion, despite the limitation that this study is primarily computational and needs to be confirmed experimentally, it demonstrates that RCC and LCC have different genomic and proteomic profiles, whereas LCC and rectal cancers are related but distinct. Ultimately, optimal management of colorectal cancers will require identifying and putting in context the relevant DNA, RNA and protein molecular signatures with respect to anatomical location and clinical behavior.

## Materials and Methods

### Somatic mutation data

We obtained Mutation Annotation Format (MAF) from TCGA somatic mutation data for 633 samples of colorectal cancers (CRCs) from the GDC legacy archive (accessed on Jan 23rd, 2018).[12] Patients demographics and clinical characteristics were obtained from cBioPortal[53,54] and Firehose (https://gdac.broadinstitute.org/, TCGA data version 2016_01_28 for COAD). CpG island methylator phenotype (CIMP) status was obtained from previously published data by Grasso et al.[55] (S22 Table). Patients were divided into non-hypermutated, hypermutated and POLE mutations positive associated CRCs. Patients with multiple sequenced samples were identified and only unique samples were retained for further analyses. Among the non-hypermutated group, 387 samples had associated clinical data and anatomical sites and were therefore retained for final analysis. These samples were divided into right-sided colon cancers, including cecum, ascending colon, hepatic flexure (n = 142), left-sided colon cancers, including splenic flexure, descending colon, sigmoid colon (n = 156) and rectal cancers (n = 89). Transverse colon and rectosigmoid tumors were excluded from the analyses. Hyper-mutated phenotypes and POLE mutated samples were also excluded from further analysis because of their low numbers to allow adequate statistical evaluation (S10 Fig). Somatic mutation analysis of non-hypermutated TCGA samples revealed RCC to have higher somatic mutations (mean = 180.5 mutations per sample) compared to LCC (mean = 132.2 mutations per sample) and rectal cancers (mean = 118.4 mutations per sample). We also obtained stage IV metastatic colorectal cancer targeted gene panel mutation data (which included 314 LCC, 200 RCC, and 189 rectal cancer samples) that has been previously published by Memorial Sloan Kettering Cancer Center (MSKCC).[11] Mutation data from MSKCC was converted into MAF files using Oncotator.[56] Hypermutated samples with associated clinical data were excluded for analysis due to their small sample size.

### Patient characteristics

Patient demographics and clinical characteristics from TCGA are shown in S23 Table. Most patents were male (54.3%). Median age at diagnosis was 66.6 years. 43.4% patients were white. Data included patients from all four stages; stage I 17.8% (n = 69), stage II 32.5% (n = 126), stage III 29.7% (n = 115) and stage IV 17.5% (n = 68). MSKCC patient demographics are described by Yaeger et al. in their original paper.[11] In MSKCC dataset, we only analyzed patients with stage IV disease (n = 703). The samples included both primary biopsy sites (n = 270) as well as metastatic biopsy sites (n = 433).

### Pipeline for Cancer Inference (PiCnIc) for evolutionary trajectories

143 LCC, 135 RCC and 76 rectal cancer samples from the TCGA data set with matched SNV/InDELs and copy number data were analyzed. We applied the Pipeline for Cancer Inference (PiCnIc) algorithm for Evolutionary Trajectory on cross-sectional TCGA data at default parameters as scripts/commands provided by authors in their R-Code (https://github.com/BIMIB-DISCo/PiCnIc-COADREAD/tree/master/scripts).[13] We interrogated 33 genes that were selected by Muzny et al. in their original report on colorectal cancer.[8] In addition, we also selected 17 unique genes from Pan Cancer analysis as reported by Ciriello et al. since these alterations are ubiquitous in solid tumors and are tissue agnostic.[57] All parameters were run on default parameters, including exclusivity among genes in the WNT (*APC* and *CTNNB1*) and RAF (*KRAS*, *NRAS*, and *BRAF*) pathways as described by Caravagna et al.[13] We ran CAPRI as the last step of PiCnIc. We carried out Mann-Withney *U* test with statistical significance 0.05, after 100 nonparametric bootstrap iterations. We considered the Akaike information criterion (AIC) score of 40% and the Bayesian information criterion (BIC) score of 30% significant. Nonparametric and statistical bootstrap estimations are shown in S1-3 Tables.

### Significantly mutated gene analysis

Consensus Driver is a relatively new driver prediction algorithm that integrates previous driver prediction algorithms (fathmm, CHASM, OncoIMPACT, DriverNet, MutSigCV, OncodriveFM) and significantly improves the quality of predictions and discovery of novel significantly mutated genes.[58] MAF files were parsed into individual sample files and converted to vcf files using MAF2VCF perl script (https://github.com/mskcc/vcf2maf) and passed through the ConsensusDriver Algorithm using colon adenocarcinoma and rectum adenocarcinoma cancer subtype options as described by Bertrand et al.[14] The resulting genes were combined into a list. Known false positive driver mutations, as previously identified by Bertrand et al.[14] were removed from our analysis. Frequencies of the identified genes were compared among each tumor location in each data set. Silent mutations were not removed from the final analysis. The enrichment analysis between RCC, LCC and rectum was carried out by chi-square and p-value <0.05, with mutational frequency ≥ 5 % considered significant.

Hotspot Analysis was carried out using the algorithm described by Chang et al.[30] R scripts publicly available were applied to our data with default parameters. MAF tools were used to create lollipop plots for drivers.[59]

### Somatic copy number alterations analysis

We obtained copy number data from firehose broad (https://gdac.broadinstitute.org/, TCGA data version 2016_01_28 for COAD) and included 164 samples of LCC, 147 samples of RCC and 99 samples of rectal cancer. Since rectal cancer samples were significantly lower compared to RCC and LCC, they were removed from further analysis as this could have led to false amplifications or deletions in RCC or LCC. Somatic copy number alterations analysis of LCC and RCC was performed by using the GISTIC algorithm on Genepattern to identify focal alterations (focal amplifications and deletions).[60,61] A q-value of 0.05 and confidence level 0.99 were considered significant. We used previously described criteria from Campbell et al. to identify significant amplifications and deletion.[62] Briefly, If GISTIC2.0 estimated the focal copy number ratio to be > 0.1 this was considered to represent focal amplification, and if it estimated the ratio to be <-0.1 then this was considered to represent deletion, respectively. Furthermore, if known target gene of each peak was the same or if peak genomic locations overlap +/-1 Megabase and if each overlapping was smaller than 10 megabase and had <25 genes then 2 peaks were considered same. Candidate genes related to significant amplifications and deletions were identified using pan-cancer patterns of somatic copy-number alteration, as described by Zack et al.[63]

### RNA sequence analysis

We identified 698 CRCs samples from the TCGA data set (COAD & READ) by using the GDC Data Portal for Cancer Genomics for detecting differential expression (accessed on January 2018).[12] Patients with multiple samples sequenced were identified and only unique samples was retained for further analyses. Normal samples were removed from the data. Again, hypermutated samples were excluded from the analyses due to small sample size, especially LCC where only 5 samples were available for analysis. Final analysis included 166 samples of left colon cancers, 146 samples of right colon cancers and 101 samples of rectal cancers.

In our analysis, LCC and rectal cancer were set as controls and were compared to RCC. In addition, LCC was set as control when differential gene expression was compared to rectal cancer. Therefore, differentially expressed genes with positive log2 value have higher expression in RCC compared to LCC and rectal cancer. Similarly, in comparing LCC and rectal cancer differentially expressed genes with positive log2 value have higher expression in rectal cancer compared to LCC.

Gene quantification are htseq-count data and were obtained from the TCGA portal. To compare gene expression between RCC and LCC, we calculated the fraction of the reads assigned to each gene relative to the total number of reads and with respect to the entire RNA repertoire which may vary drastically from sample to sample. We conducted normalization of the samples using estimateSizeFactors of the DESeq2 package.[64] This function calculates size factors for each sample using the median of ratios method. The read counts are transformed to the log scale. DESeq analysis yielded the base means across samples, log2 fold changes, standard errors, test statistics, p-values and adjusted p-values. A p-value ≤ 0.05 was used as a cut off to signify statistical significance.

For our analysis, a cut off value of log2 fold change ≤ or ≥ ±1.5 and adjusted p ≤ 0.05 was used to identify differentially expressed genes (DEG). Downregulated genes are defined as log2 fold change ≤ −1.5 and adjusted p ≤ 0.05; upregulated genes are defined as log2 fold change ≥ +1.5 and adjusted p ≤ 0.05. Using the widely used current version of The Database for Annotation, Visualization and Integrated Discovery (DAVID 6.8) functional annotation clustering[36,37] was applied to our differentially expressed genes between RCC vs LCC, RCC vs rectal cancers and LCC vs rectal cancers. We used the high stringency filter and clusters with p-value <0.05 were considered significant.

We also employed a hierarchical (agglomerative) clustering tool, REVIGO[38] using gene ontology (GO) terms for biological processes enriched in our data from DAVID to identify differences and similarities between RCC, LCC and rectal cancers. A p-value of <0.05 was considered significant.

Using Search Tool for the Retrieval of Interacting Genes (STRING)[65] data on protein-protein interactions (PPI) with minimum required interaction score of 0.04 were obtained. We employed the novel graph theoretic clustering algorithm, Molecular Complex Detection (MCODE), to identify the most significant module in the PPI network[39]. The enrichment analysis of the DEG involved in the modules was also carried out with DAVID. Pathway and network enrichment analysis of p-value <0.05 were considered significant.

### MicroRNA analysis

551 miRNA CRC samples from the TCGA colon and rectal cancer data base (COAD & READ) were analyzed using the GDC Data Portal for Cancer Genomics (accessed on February,2018).[12] Samples with POLE gene mutations and MSI-H were excluded because of their small numbers. Final analysis included 147 samples of LCC, 135 samples of RCC and 90 samples of rectal cancer. To compare microRNA expression patterns in RCC, LCC and rectal cancer, similar methodology as described above for RNA sequence analysis was used.[64] A cutoff of log2FC > 0.584 (Log2FC>0.584 = FC>1.5) and adjusted p <0.05 were defined as significant. Further network analysis was carried out using a newer version, ONCO.IO (https://onco.io), of the MicroRNA OncoBase (MirOB) web tool (http://mirob.interactome.ru/). Significant candidate microRNAs were analyzed to find key target genes, transcription factors, pathological processes and diseases associated with these miRNAs. MiRNAs shared between RCC, LCC and rectal cancers were used to create a network as shown in S19 Table. Similarly, uniquely dysregulated miRNAs between RCC vs LCC and RCC vs rectal cancers were used to build networks, as shown in S20 and S21 Tables.

### Proteomics analysis

Reverse Phase Protein Array (RPPA) samples (COAD & READ) were downloaded from The Cancer Proteome Atlas (TCPA) website (http://tcpaportal.org/tcpa/).[40] The samples included 109 left-sided colon cancer, 98 right-sided colon cancer and 67 rectal cancer samples. Proteomics analysis was performed by using our previously described methods.[41]

As shown in S24 Table, the rankings of the association estimators identified by the average performance score (Avg) varied according to the constructed module sizes and the selected groups. In the pathway-level analysis, best performing methods based on the selected groups are as follows: MM for RCC, KDE for LCC and rectal cancers (S24a Table). Similarly, in the gene-level analysis, best performing methods based on the selected groups are as follows: MM for RCC, KDE for LCC and rectal cancers (S24b Table).

The sub networks of the module hub genes identified by the association estimators, which provide the highest precision scores based on a statistical test (with a p-value <0.05), are generated by Cytoscape[66] for all selected groups (S25 Table). The genes registered in DisGeNET[67] and experimentally confirmed for the diseases, are shown in Figs 7a-c with colored and larger nodes. Among those, genes that are not colored but have a red frame have a PMID value of one. There is no entry in DisGeNET for the grey colored nodes. Also, the most associated top three biological pathways, to which each module is related, are given above or below the relevant module to annotate each module.

## Supporting information

S1 Fig shows clonal evolution in Colorectal cancer using CAPRI algorithm. The events of the model are connected by dashed lines where red dotted lines denote hard and orange denotes soft exclusivity. Algorithm uses both Bayesian information criterion ‘BIC’ and Akaike information criterion ‘AIC’ as a regularization. Non-parametric bootstrap scores (NPB) are shown in the figure with hypergeometric test p-value cutoff of <0.05. Other relations including temporal priority, probability raising are shown in Figs 1a-c and reported data in S1-3 Tables. 1a) clonal evolution in RCC, 1b) clonal evolution in LCC and 1c) clonal evolution in rectal cancers.

S2 Fig. Lollipop plots of driver genes enriched in RCC compared to LCC and rectal cancers.

S3 Fig. Novel genes discovered in our analysis in RCC showing tendency towards mutual exclusivity.

S4a Fig. Significantly mutated genes from primary tumor site biopsy and metastatic site biopsy in right-sided colon cancer

S4b Fig. Significantly mutated genes from primary tumor site biopsy and metastatic site biopsy in rectal cancers

S5a Fig. Oncoplot showing mutual exclusivity of AMER1 and other keys genes of β-Catenin destruction complex in RCC.

S5b Fig. Oncoplot showing mutual exclusivity of AMER1 and other keys genes of β-Catenin destruction complex in LCC.

S5c Fig. Oncoplot showing mutual exclusivity of AMER1 and other keys genes of β-Catenin destruction complex in rectal cancer.

S6a Fig. Copy number alterations in left-sided colon cancers showing amplified regions (red) and deleted regions (blue)

S6b Fig. Copy number alterations in right-sided colon cancers showing amplified regions (red) and deleted regions (blue)

S7 Fig. Dysregulated biological processes between a) right-sided colon cancer and left-sided colon cancer and b) right-sided colon cancer and rectal cancers. Red bars show similar biological processes and blue bars show different biological processes between RCC/LCC vs RCC/rectal cancers.

S8a Fig. Interactive view of dysregulated biological processes between right colon cancers and left colon cancers using REVIGO. Key pathways include cell-differentiation, negative regulation of inflammatory response, embryonic limb development, epidermis development and lipoprotein biosynthesis.

S8b Fig. Interactive view of dysregulated biological processes between right colon cancers and rectal cancers using REVIGO (using both upregulated and downregulated GO terms)

S8c Fig. Interactive view of dysregulated biological processes between left colon cancers and rectal cancers using REVIGO (using both upregulated and downregulated GO terms)

S9a Fig shows a network of microRNAs, their transcription factors and target genes which are shared between RCC vs LCC and RCC vs rectal cancers. (diamond, magenta color) = microRNAs, circles = target genes, triangle (orange color) = transcription factors, shape = receptor.

S9b Fig shows unique microRNAs and their target genes between RCC and LCC. (excluding shared micro RNAs between RCC/LCC and RCC/rectal cancers).

S9c Fig shows unique microRNAs and their target genes between RCC and rectal cancers.

S10 Fig. Inclusion and exclusion criteria for somatic mutation analysis. MSI-H, POLE mutation samples, rectosigmoid and transverse colon cancers were excluded (highlighted green).

S1 Table. Right colon cancer (RCC) PicNiC statistics (bic and aic)

S2 Table. Left colon cancer (LCC) PicNiC statistics (bic and aic)

S3 Table. Rectal cancer PicNiC statistics (bic and aic)

S4 Table. Genes from Pan Cancer analyses by Ciriello et al. and TCGA Comprehensive Molecular Characterization of Human Colon and Rectal Cancer (2012)

S5 Table. Enriched consensusDriver (highlighted light green) between LCC vs RCC, LCC vs rectal cancer and RCC vs rectal cancer - TCGA

S6 Table. Enriched consensusDriver (highlighted light green) between LCC vs RCC, LCC vs rectal cancer and RCC vs rectal cancer – MSKCC

S7 Table. Enriched consensusDriver (highlighted light green) between Primary tumor site biopsy (stage IV) vs metastatic site biopsy – MSKCC

S8 Table. Amplified and deleted regions along with candidate genes between RCC and LCC

S9 Table. Downregulated and upregulated genes between RCC and LCC_Padj <0.05 Log2FC>1.5

S10 Table. Downregulated and upregulated genes between RCC and rectal cacner_padj<0.05 Log2FC>1.5

S11 Table. Downregulated and upregulated genes between LCC and rectal cancer_Padj<0.05 Log2FC>1.5

S12 Table. Functional annotation clustering of upregulated and downregulated genes (RCC vs LCC log2FC greater than 1.5 and padj less than 0.05) using DAVID _ high stringency filter

S13 Table. Functional annotation clustering of upregulated and downregulated genes (RCC vs rectal cancer log2FC greater than 1.5 and padj less than 0.05) using DAVID _ high stringency filter

S14 Table. Functional annotation clustering of upregulated and downregulated genes (LCC vs rectal cancer log2FC greater than 1.5 and padj less than 0.05) using DAVID _ high stringency filter

S15 Table. Dysregulated biological processes RCC vs LCC-REVIGO (upregulated and downregulated GO terms using REVIGO with medium similarity filter)

S16 Table. Dysregulated biological processes RCC vs Rectal cancers-REVIGO (upregulated and downregulated GO terms using REVIGO with medium similarity filter)

S17 Table. Dysregulated biological processes LCC vs Rectal cancers-REVIGO (upregulated and downregulated GO terms using REVIGO with medium similarity filter)

S18 Table. Upregulated and downregulated microRNAs between RCC vs LCC, RCC vs rectal cancers and LCC vs rectal cancers (MSS)

S19 Table. Dysregulated microRNAs shared between RCC, LCC and rectal cancers with their transcription factors, target genes, pathological process and diseases.

S20 Table. Uniquely dysregulated microRNAs between RCC and LCC with their transcription factors, target genes, pathological process and diseases

S21 Table. Uniquely dysregulated microRNAs between RCC and rectal cancers with their transcription factors, target genes, pathological process and diseases.

S22 Table. CpG island methylator phenotype (CIMP) status data with excluded samples

S23 Table. Patients demographics from TCGA

S24 Table. Pathway and gene level precision ratios by module numbers for each selected group (P-value <0.05).

S25 Table. Hub genes identified by the association estimators with the highest precision scores with a p-value <0.05 for right-sided colon cancer, left-sided colon cancer and rectal cancers

## Acknowledgements

Part of A.H.’s time was supported by a Merit Review Award (I01 BX000545), Medical Research Service, Department of Veterans Affairs

## References

1. Lee GH, Malietzis G, Askari A, Bernardo D, Al-Hassi HO, Clark SK. Is right-sided colon cancer different to left-sided colorectal cancer? - A systematic review. Eur J Surg Oncol. Elsevier Ltd; 2015;41: 300–308. doi:10.1016/j.ejso.2014.11.001

2. Meguid RA, Slidell MB, Wolfgang CL, Chang DC, Ahuja N. Is there a difference in survival between right- versus left-sided colon cancers? Ann Surg Oncol. 2008;15: 2388–2394. doi:10.1245/s10434-008-0015-y

3. van der Sijp MPL, Bastiaannet E, Mesker WE, van der Geest LGM, Breugom AJ, Steup WH, et al. Differences between colon and rectal cancer in complications, short-term survival and recurrences. Int J Colorectal Dis. 2016;31: 1683–1691. doi:10.1007/s00384-016-2633-3

4. Venook AP, Niedzwiecki D, Lenz H-J, Innocenti F, Mahoney MR, O’Neil BH, et al. CALGB/SWOG 80405: Phase III trial of irinotecan/5-FU/leucovorin (FOLFIRI) or oxaliplatin/5-FU/leucovorin (mFOLFOX6) with bevacizumab (BV) or cetuximab (CET) for patients (pts) with KRAS wild-type (wt) untreated metastatic adenocarcinoma of the colon or re. J Clin Oncol. 2014;32: LBA3–LBA3. doi:10.1200/jco.2014.32.18_suppl.lba3

5. Heinemann V, von Weikersthal LF, Decker T, Kiani A, Vehling-Kaiser U, Al-Batran S-E, et al. FOLFIRI plus cetuximab versus FOLFIRI plus bevacizumab as first-line treatment for patients with metastatic colorectal cancer (FIRE-3): a randomised, open-label, phase 3 trial. Lancet Oncol. 2014;15: 1065–1075. doi:10.1016/S1470-2045(14)70330-4

6. Sanz-Pamplona R, Cordero D, Berenguer A, Lejbkowicz F, Rennert H, Salazar R, et al. Gene Expression Differences between Colon and Rectum Tumors. Clin Cancer Res. 2011;17: 7303–7312. doi:10.1158/1078-0432.CCR-11-1570

7. Hong TS, Clark JW, Haigis KM. Cancers of the Colon and Rectum: Identical or Fraternal Twins? Cancer Discov. 2012;2: 117–121. doi:10.1158/2159-8290.CD-11-0315

8. Muzny DM, Bainbridge MN, Chang K, Dinh HH, Drummond JA, Fowler G, et al.Comprehensive molecular characterization of human colon and rectal cancer. Nature. Nature Publishing Group; 2012;487: 330–337. doi:10.1038/nature11252

9. Yu J, Wu WKK, Li X, He J, Li XX, Ng SSM, et al. Novel recurrently mutated genes and a prognostic mutation signature in colorectal cancer. Gut. 2015;64: 636–645. doi:10.1136/gutjnl-2013-306620

10. Hu W, Yang Y, Li X, Huang M, Xu F, Ge W, et al. Multi-omics Approach Reveals Distinct Differences in Left- and Right-sided Colon Cancer. Mol Cancer Res. 2017; molcanres.0483.2017. doi:10.1158/1541-7786.MCR-17-0483

11. Yaeger R, Chatila WK, Lipsyc MD, Hechtman JF, Cercek A, Sanchez-Vega F, et al. Clinical Sequencing Defines the Genomic Landscape of Metastatic Colorectal Cancer. Cancer Cell. Elsevier Inc.; 2018;33: 125–136.e3. doi:10.1016/j.ccell.2017.12.004

12. Grossman RL, Heath AP, Ferretti V, Varmus HE, Lowy DR, Kibbe WA, et al. Toward a Shared Vision for Cancer Genomic Data. N Engl J Med. 2016;375: 1109–12. doi:10.1056/NEJMp1607591

13. Caravagna G, Graudenzi A, Ramazzotti D, Sanz-Pamplona R, De Sano L, Mauri G, et al. Algorithmic methods to infer the evolutionary trajectories in cancer progression. Proc Natl Acad Sci. 2016;113: E4025–E4034. doi:10.1073/pnas.1520213113

14. Bertrand D, Drissler S, Chia B, Koh JY, Li C, Suphavilai C, et al. ConsensusDriver Improves Upon Individual Algorithms For Predicting Driver Alterations In Different Cancer Types And Individual Patients — A Toolbox For Precision Oncology. bioRxiv. 2017;

15. Vogelstein B, Papadopoulos N, Velculescu VE, Zhou S, Diaz LA, Kinzler KW. Cancer genome landscapes. Science. 2013;339: 1546–58. doi:10.1126/science.1235122

16. Jamal-Hanjani M, Wilson GA, McGranahan N, Birkbak NJ, Watkins TBK, Veeriah S, et al. Tracking the Evolution of Non-Small-Cell Lung Cancer. N Engl J Med. 2017;376: 2109–2121. doi:10.1056/NEJMoa1616288

17. Phillips PC. Epistasis—the essential role of gene interactions in the structure and evolution of genetic systems. Nat Rev Genet. 2008;9: 855–867. doi:10.1038/nrg2452.Epistasis

18. Ortmann CA, Kent DG, Nangalia J, Silber Y, Wedge DC, Grinfeld J, et al. Effect of Mutation Order on Myeloproliferative Neoplasms. N Engl J Med. 2015;372: 601–612. doi:10.1056/NEJMoa1412098

19. Haan JC, Labots M, Rausch C, Koopman M, Tol J, Mekenkamp LJM, et al. Genomic landscape of metastatic colorectal cancer. Nat Commun. Nature Publishing Group; 2014;5: 1–12. doi:10.1038/ncomms6457

20. Tian Z-Q, Shi J-W, Wang X-R, Li Z, Wang G-Y. New cancer suppressor gene for colorectal adenocarcinoma: Filamin A. World J Gastroenterol. 2015;21: 2199–2205. doi:10.3748/wjg.v21.i7.2199

21. Nagai T, Mizuno K. Multifaceted roles of Furry proteins in invertebrates and vertebrates. J Biochem. 2014;155: 137–146. doi:10.1093/jb/mvu001

22. Sato N, Fukushima N, Chang R, Matsubayashi H, Goggins M. Differential and Epigenetic Gene Expression Profiling Identifies Frequent Disruption of the RELN Pathway in Pancreatic Cancers. Gastroenterology. 2006;130: 548–565. doi:10.1053/j.gastro.2005.11.008

23. Soltani MH, Pichardo R, Song Z, Sangha N, Camacho F, Satyamoorthy K, et al. Microtubule-Associated Protein 2, a Marker of Neuronal Differentiation, Induces Mitotic Defects, Inhibits Growth of Melanoma Cells, and Predicts Metastatic Potential of Cutaneous Melanoma. Am J Pathol. 2005;166: 1841–1850. doi:10.1016/S0002-9440(10)62493-5

24. Crnic I, Strittmatter K, Cavallaro U, Kopfstein L, Jussila L, Alitalo K, et al. Loss of Neural Cell Adhesion Molecule Induces Tumor Metastasis by Up-regulating Lymphangiogenesis. Cancer Res. 2004;64: 8630–8638. doi:10.1158/0008-5472.CAN-04-2523

25. Alfayez M, Vishnubalaji R, Alajez NM. Runt-related Transcription Factor 1 (RUNX1T1) Suppresses Colorectal Cancer Cells Through Regulation of Cell Proliferation and Chemotherapeutic Drug Resistance. Anticancer Res. 2016;36: 5257–5263. doi:10.21873/anticanres.11096

26. Jethwa A, Słabicki M, Hüllein J, Jentzsch M, Dalal V, Rabe S, et al. TRRAP is essential for regulating the accumulation of mutant and wild-type p53 in lymphoma. Blood. 2018; doi:10.1182/blood-2017-09-806679

27. Ying J, Li H, Seng TJ, Langford C, Srivastava G, Tsao SW, et al. Functional epigenetics identifies a protocadherin PCDH10 as a candidate tumor suppressor for nasopharyngeal, esophageal and multiple other carcinomas with frequent methylation. Oncogene. 2006;25: 1070–1080. doi:10.1038/sj.onc.1209154

28. Wang Z, Sun P, Gao C, Chen J, Li J, Chen Z, et al. Down-regulation of LRP1B in colon cancer promoted the growth and migration of cancer cells. Exp Cell Res. 2017;357: 1–8. doi:10.1016/j.yexcr.2017.04.010

29. Sanz-Pamplona R, Lopez-Doriga A, Paré-Brunet L, Lázaro K, Bellido F, Alonso MH, et al. Exome sequencing reveals AMER1 as a frequently mutated gene in colorectal cancer. Clin Cancer Res. 2015;21: 4709–4718. doi:10.1158/1078-0432.CCR-15-0159

30. Chang MT, Asthana S, Gao SP, Lee BH, Chapman JS, Kandoth C, et al. Identifying recurrent mutations in cancer reveals widespread lineage diversity and mutational specificity. Nat Biotechnol. 2016;34: 155–63. doi:10.1038/nbt.3391.

31. Imperial R, Toor OM, Hussain A, Subramanian J, Masood A. Comprehensive pancancer genomic analysis reveals (RTK)-RAS-RAF-MEK as a key dysregulated pathway in cancer: Its clinical implications. Semin Cancer Biol. 2017; doi:10.1016/j.semcancer.2017.11.016

32. Narayan S, Roy D. Role of APC and DNA mismatch repair genes in the development of colorectal cancers. Mol Cancer. 2003;2: 41. doi:10.1186/1476-4598-2-41

33. Miyoshi Y, Nagase H, Ando H, Horii A, Ichii S, Nakatsuru S, et al. Somatic mutations of the APC gene in colorectal tumors: mutation cluster region in the APC gene. Hum Mol Genet. 1992;1: 229–33.

34. Christie M, Jorissen RN, Mouradov D, Sakthianandeswaren A, Li S, Day F, et al. Different APC genotypes in proximal and distal sporadic colorectal cancers suggest distinct WNT/ β -catenin signalling thresholds for tumourigenesis. Oncogene. 2013;32: 4675–4682. doi:10.1038/onc.2012.486

35. Forbes SA, Beare D, Boutselakis H, Bamford S, Bindal N, Tate J, et al. COSMIC: somatic cancer genetics at high-resolution. Nucleic Acids Res. 2017;45: D777–D783. doi:10.1093/nar/gkw1121

36. Huang DW, Sherman BT, Lempicki RA. Systematic and integrative analysis of large gene lists using DAVID bioinformatics resources. Nat Protoc. 2009;4: 44–57. doi:10.1038/nprot.2008.211

37. Huang DW, Sherman BT, Lempicki RA. Bioinformatics enrichment tools: Paths toward the comprehensive functional analysis of large gene lists. Nucleic Acids Res. 2009;37: 1–13. doi:10.1093/nar/gkn923

38. Supek F, Bošnjak M, Škunca N, Šmuc T. Revigo summarizes and visualizes long lists of gene ontology terms. PLoS One. 2011;6: e21800. doi:10.1371/journal.pone.0021800

39. Bader GD, Hogue CW. An automated method for finding molecular complexes in large protein interaction networks. BMC Bioinformatics. 2003;4: 2. doi:10.1186/1471-2105-4-2

40. Li J, Lu Y, Akbani R, Ju Z, Roebuck PL, Liu W, et al. TCPA: a resource for cancer functional proteomics data. Nat Methods. 2013;10: 1046–1047. doi:10.1038/nmeth.2650

41. Erdoğan C, Kurt Z, Diri B. Estimation of the proteomic cancer co-expression sub networks by using association estimators. Yamanishi Y, editor. PLoS One. 2017;12: e0188016. doi:10.1371/journal.pone.0188016

42. Luo Y, Wang X, Wang H, Xu Y, Wen Q, Fan S, et al. High Bak Expression Is Associated with a Favorable Prognosis in Breast Cancer and Sensitizes Breast Cancer Cells to Paclitaxel. Castresana JS, editor. PLoS One. 2015;10: e0138955. doi:10.1371/journal.pone.0138955

43. Tang J, Xi S, Wang G, Wang B, Yan S, Wu Y, et al. Prognostic significance of BRCA1-associated protein 1 in colorectal cancer. Med Oncol. 2013;30: 541. doi:10.1007/s12032-013-0541-8

44. Greig FH, Nixon GF. Phosphoprotein enriched in astrocytes (PEA)-15: A potential therapeutic target in multiple disease states. Pharmacol Ther. 2014;143: 265–274. doi:10.1016/j.pharmthera.2014.03.006

45. Zhang L, Kim SB, Luitel K, Shay JW. Cholesterol Depletion by TASIN-1 Induces Apoptotic Cell Death through the ER Stress/ROS/JNK Signaling in Colon Cancer Cells. Mol Cancer Ther. 2018; molcanther.0887.2017. doi:10.1158/1535-7163.MCT-17-0887

46. Zhang L, Theodoropoulos PC, Eskiocak U, Wang W, Moon Y-A, Posner B, et al. Selective targeting of mutant adenomatous polyposis coli (APC) in colorectal cancer. Sci Transl Med. 2016;8: 361ra140 LP–361ra140.

47. Vignot S, Besse B, André F, Spano J-P, Soria J-C. Discrepancies between primary tumor and metastasis: A literature review on clinically established biomarkers. Crit Rev Oncol Hematol. 2012;84: 301–313. doi:10.1016/j.critrevonc.2012.05.002

48. Bhatlekar S, Fields JZ, Boman BM. HOX genes and their role in the development of human cancers. Journal of Molecular Medicine. 2014. pp. 811–823. doi:10.1007/s00109-014-1181-y

49. Luo X, Cheng C, Tan Z, Li N, Tang M, Yang L, et al. Emerging roles of lipid metabolism in cancer metastasis. Mol Cancer. Molecular Cancer; 2017;16: 1–10. doi:10.1186/s12943-017-0646-3

50. Bar-Shavit R, Maoz M, Kancharla A, Nag JK, Agranovich D, Grisaru-Granovsky S, et al. G protein-coupled receptors in cancer. Int J Mol Sci. 2016;17: 1–16. doi:10.3390/ijms17081320

51. Williams GH, Stoeber K. The cell cycle and cancer. J Pathol. 2012;226: 352–364. doi:10.1002/path.3022

52. Jingwen B, Yaochen L, Guojun Z. Cell cycle regulation and anticancer drug discovery. Cancer Biol Med. 2017;14: 348. doi:10.20892/j.issn.2095-3941.2017.0033

53. Gao J, Aksoy BA, Dogrusoz U, Dresdner G, Gross B, Sumer SO, et al. Integrative Analysis of Complex Cancer Genomics and Clinical Profiles Using the cBioPortal. Sci Signal. 2013;6: 1–34. doi:10.1126/scisignal.2004088.Integrative

54. Cerami E, Gao J, Dogrusoz U, Gross BE, Sumer SO, Aksoy BA, et al. The cBio Cancer Genomics Portal: An open platform for exploring multidimensional cancer genomics data. Cancer Discov. 2012;2: 401–404. doi:10.1158/2159-8290.CD-12-0095

55. Grasso CS, Giannakis M, Wells DK, Hamada T, Mu XJ, Quist M, et al. Genetic mechanisms of immune evasion in colorectal cancer. Cancer Discov. 2018; CD-17-1327. doi:10.1158/2159-8290.CD-17-1327

56. Ramos AH, Lichtenstein L, Gupta M, Lawrence MS, Pugh TJ, Saksena G, et al. Oncotator: Cancer variant annotation tool. Hum Mutat. 2015;36: E2423–E2429. doi:10.1002/humu.22771

57. Ciriello G, Miller ML, Aksoy BA, Senbabaoglu Y, Schultz N, Sander C. Emerging landscape of oncogenic signatures across human cancers. Nat Genet. Nature Publishing Group; 2013;45: 1127–1133. doi:10.1038/ng.2762

58. Alexandrov LB, Nik-Zainal S, Wedge DC, Aparicio SAJR, Behjati S, Biankin A V., et al. Signatures of mutational processes in human cancer. Nature. 2013;500: 415–421. doi:10.1038/nature12477

59. Mayakonda A, Koeffler HP. Maftools: Efficient analysis, visualization and summarization of MAF files from large-scale cohort based cancer studies. bioRxiv. 2016; 052662. doi:10.1101/052662

60. Mermel CH, Schumacher SE, Hill B, Meyerson ML, Beroukhim R, Getz G. GISTIC2.0 facilitates sensitive and confident localization of the targets of focal somatic copy-number alteration in human cancers. Genome Biol. 2011;12: 1–14. doi:10.1186/gb-2011-12-4-r41

61. Reich M, Liefeld T, Gould J, Lerner J, Tamayo P, Mesirov JP. GenePattern 2.0. Nat Genet. 2006;38: 500–501.

62. Campbell JD, Alexandrov A, Kim J, Wala J, Berger AH, Pedamallu CS, et al. Distinct patterns of somatic genome alterations in lung adenocarcinomas and squamous cell carcinomas. Nat Genet. 2016;48: 607–616.

63. Zack TI, Schumacher SE, Carter SL, Cherniack AD, Saksena G, Tabak B, et al. Pan-cancer patterns of somatic copy-number alteration. Nat Genet. 2013;45: 1134–1140. doi:1038/ng.2760. Pan-cancer

64. Love MI, Huber W, Anders S. Moderated estimation of fold change and dispersion for RNA-seq data with DESeq2. Genome Biol. 2014;15: 1–21. doi:10.1186/s13059-014-0550-8

65. Szklarczyk D, Morris JH, Cook H, Kuhn M, Wyder S, Simonovic M, et al. The STRING database in 2017: quality-controlled protein – protein association networks, made broadly accessible. Nucleic Acids Res. 2017;45: 362–368. doi:10.1093/nar/gkw937

66. Shannon P, Markiel A, Owen Ozier 2, Baliga NS, Wang JT, Ramage D, et al.Cytoscape: a software environment for integrated models of biomolecular interaction networks. Genome Res. 2003; 2498–2504. doi:10.1101/gr.1239303.metabolite

67. Piñero J, Queralt-Rosinach N, Bravo À, Deu-Pons J, Bauer-Mehren A, Baron M, et al. DisGeNET: A discovery platform for the dynamical exploration of human diseases and their genes. Database. 2015;2015: 1–17. doi:10.1093/database/bav028

